# RNA binding protein RBM3 augments kissing loop formation with lncRNAs to enhance translational control

**DOI:** 10.1101/2021.12.14.472669

**Authors:** Afreen Asif Ali Sayed, Sonali Choudhury, Dharmalingam Subramaniam, Sumedha Gunewardena, Sivapriya Ponnurangam, Prasad Dandawate, Ossama Tawfik, David Standing, Subhash B. Padhye, Linheng Li, Tomoo Iwakuma, Shahid Umar, Roy A. Jensen, Sufi Mary Thomas, Shrikant Anant

## Abstract

It is becoming apparent that translational regulation involves the coordinated actions of RNA binding proteins (RBPs) and non-coding RNAs. For efficient translation, mRNA needs to be circularized, which is catalyzed by RNA binding proteins and translation factors. However, the role of lncRNAs in the process is not yet defined. We first performed RNA-seq and RNA-immunoprecipitation coupled-Seq and identified LSAMP-3 and Flii-1. Moreover, modeling studies suggest enhanced kissing loop interactions including of transcripts that encode angiogenesis and epithelial mesenchymal transition. While intestinal epithelial cell specific RBM3 transgenic mice showed increased LSAMP-3 and Flii-1, this was reduced in knockout mice. Also, RBM3 overexpression increased tumor xenograft growth, this was suppressed by knockdown of the lncRNAs. Also, knockdown of endogenous RBM3 reduced lncRNA levels and tumor xenograft growth. In addition, it reduced colitis-associated cancers. We propose that RBPs such as RBM3 mediate their function through regulatory lncRNAs that enable circularization to control translation.

## Introduction

RNA binding proteins (RBPs) form dynamic complexes with RNA and other proteins to regulate its fate and function^1, 2^. These ribonucleoprotein (RNP) complexes consisting of both protein-RNA and protein-protein interactions help regulate various aspects of RNA structure and function that includes mRNA translation^1, 3–5^. Any perturbation to these interactions could lead to drastic changes in gene expression which are linked to various diseases^6, 7^. Due to their central roles in the posttranscriptional gene expression, dysregulation of various RBPs have been implicated in different cancers ^6, 8^. We and others have demonstrated that RBPs including RBM3 play an important role in injury, maintenance of intestinal homeostasis, and oncogenesis^9–12^.

RBM3 is up-regulated under stress conditions such as cold shock and hypoxia ^13, 14^. The protein binds AU-rich sequences in the 3’untranslated regions (3’UTR) of rapidly degraded mRNAs such as vascular endothelial growth factor (VEGF) and interleukin8 (IL-8) and increases the stability and translation of these mRNAs^13, 15^. RBM3 also enhances global protein translation by interacting with the 60S ribosomal subunit, and transcriptionally active polysomes^16^. Further, RBM3 acts as a proto-oncogene mediating cell transformation and is upregulated in multiple solid tumors including colorectal, breast, pancreas, lung, ovary and prostate cancers^12, 16–18^. In previous studies, our laboratory established the role of RBM3 in colon cancer stem cells (CSCs) and showed that RBM3 overexpression results in increased side population and spheroid formation^19^. CSCs are a small population of cells in the tumor with self-renewal capacities contributing to tumor relapse and metastasis.

Recent advances in sequencing have drawn attention to interactions between RBPs and long non-coding RNAs (lncRNA), an important class of regulatory non-coding RNAs that have limited protein-coding potential. They generally contain more than 200 nucleotides and are transcribed by RNA polymerase II^20, 21^. LncRNAs can interact with DNA, RNA or protein giving them plasticity to act as guides, decoy, scaffolds, and signaling molecules^22, 23^. Functionally, lncRNA can regulate gene expression either transcriptionally or post-transcriptionally^24^. The role of lncRNA in regulating gene expression via chromatin remodeling and modification, binding to transcription factors, histone-modifying complexes, proteins, and RNA polymerase II, has been studied extensively^25, 26^. Further, lncRNA can also play an important role in miRNA regulation, RNA turnover and translation control^27, 28^. LncRNAs have been implicated in various functions including development, differentiation, aging, stemness, and disease. As such, few studies have examined the regulation of lncRNA by RBPs in cancer. In this study, we describe the molecular mechanisms whereby RBM3-regulated lncRNAs contributes to epithelial mesenchymal transition to facilitate colon cancer progression.

## Results

### RBM3 is overexpressed in Colon Cancer

We first examined levels of RBM3 in relation to patients and tumor characteristics in The Cancer Genome Atlas (TCGA) database. RBM3 mRNA levels were significantly higher in colon tumor samples (n=286) as compared to normal colon (n=41) (p<0.001) (Figure 1A). Transcript levels were higher in all stages of cancer compared to normal colon (
Figure S1A). Moreover, expression in colon cancer tissues were increased irrespective of age, gender, or histology (Figure S1B-D). To confirm the increase, we analyzed a cDNA array panel containing colon cancer and matched adjacent normal colon tissue samples. There were significantly higher RBM3 mRNA levels in the colon tumor samples (n=24) compared to adjacent normal colon tissue (p=0.001) (Figure 1B and Table 1). Furthermore, RBM3 protein levels were observed to be higher in a tissue microarray that contained primary adenocarcinoma, metastatic tumor and normal tissue (Figure 1C). The composite score showed significantly higher expression of RBM3 in the tumor (n=28, p=0.002) and metastasis (n=30, p=0.036) as compared to normal tissue (n=6). RBM3 expression was also increased in Stage I/II (n=15, p=0.017), Stage III/IV (n=12, p=0.041) and metastasis (n=30, p=0.036) as compared to normal (n=6) and Stage I (n=3) tumors (Figure 1D). Moreover, RBM3 protein levels were higher in the cancers irrespective of age, gender, or histology (Table 2). Additionally, western blot analysis showed increased levels of RBM3 protein in established colon cancer cell lines as compared to human normal colon epithelial FHC cell lines (Figure 1E). These data establish that RBM3 expression is upregulated in a stage-dependent manner in colon cancer.

**Figure 1:**
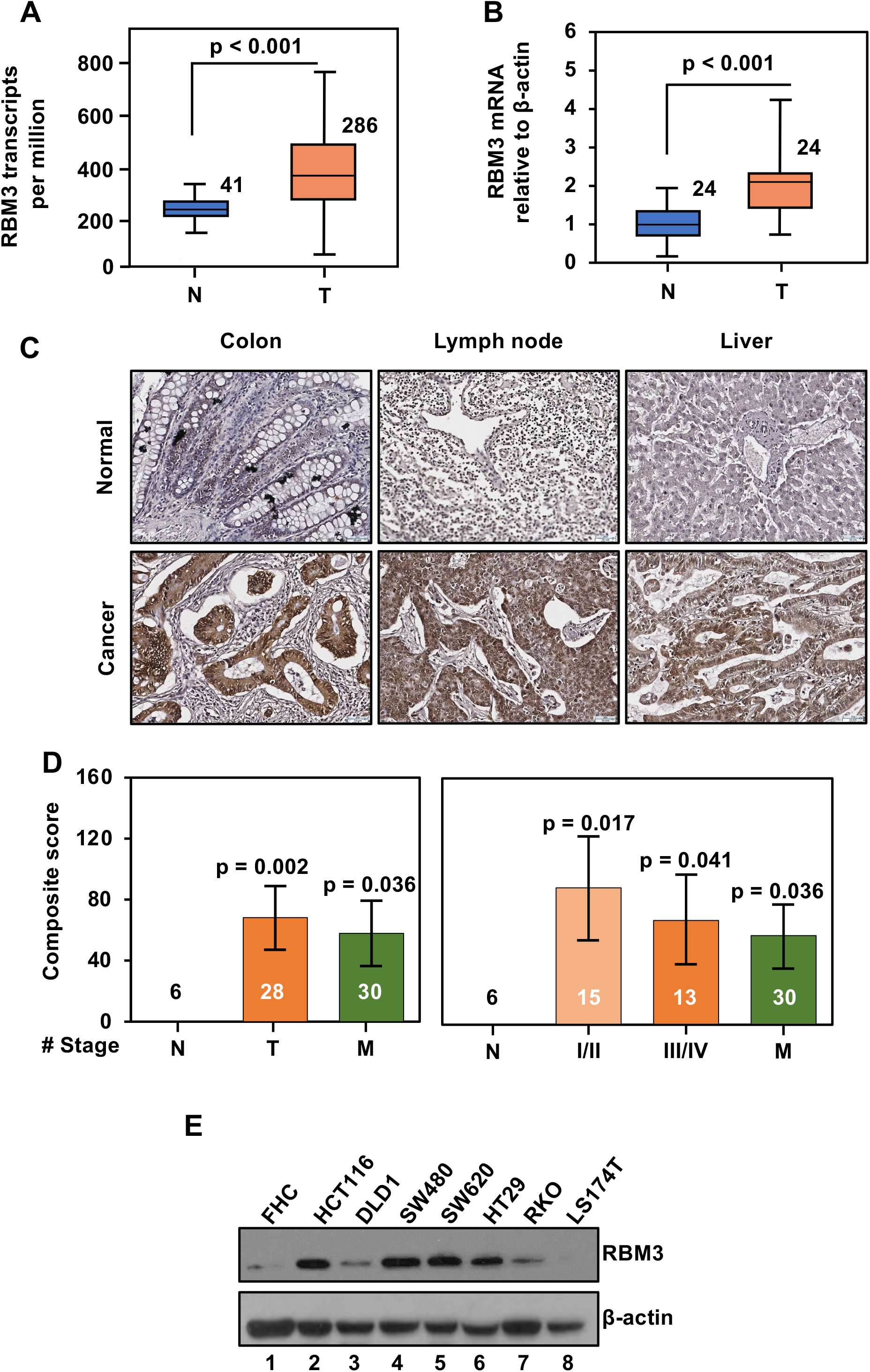
RBM3 is upregulated in Colon Cancer. (A) The Cancer Genome Atlas (TCGA) database analyses demonstrate RBM3 mRNA levels are significantly higher in colon cancer (T) (n=286) than in normal colon (N) (n=41) (p<0.001). (B) RBM3 mRNA expression from a cDNA array of colon tumor (T) samples (n=24) and matched adjacent normal colon tissue (N) (n=24) normalized to β-actin, shows a significant increase of RBM3 expression in tumor tissue (p=0.023). (C) Immunohistochemistry (IHC) of a colon cancer tumor microarray ^59^ shows that RBM3 is upregulated in colon adenocarcinoma along with lymph node metastasis and liver metastasis as compared to the normal colon, lymph node, and liver. (D) Composite score of colon cancer TMA shows significantly higher expression of RBM3 in tumor (n=28) (p=0.002) and metastasis (n=30) (p=0.036) as compared to normal tissue. RBM3 expression is also increased in the different stages of colon cancer (Stage I (n = 3), Stage II (n = 12) (p=0.015), Stage III (n= 11) (p=0.07), Stage IV (n = 2) and metastasis (n=30) (p=0.036)) as compared to normal colon, liver and lymph node (n=3 each). (E) Western blot analysis of RBM3 protein expression in established colon cancer cell lines HCT116, DLD1, SW480, SW620, HT29, RKO and LST17T as compared to normal colon epithelial cells (FHC cell line). Data in Figure 1 are represented as ± SEM. Also, see Supplementary figure 1

### RBM3 regulates long non-coding RNA expression

Upon binding to the 3’UTR of client mRNA, RBM3 affects the stability and translation of the mRNA^15, 29^. Previous studies have also shown that RBM3 regulates the posttranscriptional biogenesis of miRNA by regulating their association with dicer complexes^16^. However, the effect of RBM3 on other non-coding RNA has not been explored. Accordingly, we overexpressed RBM3 in three colon cancer cell lines with low to moderate expression of RBM3 (RKO, HCT116, and DLD1) (Figure S2A). Subsequently, we performed RNA-sequencing (RNA-seq) of HCT116 and DLD1 cell lines with RBM3 overexpression. We identified the differentially expressed lncRNA between RBM3 overexpressing and empty vector cells and visualized them by volcano plots. While RBM3-overexpressing HCT116 cells had 718 upregulated and 715 downregulated lncRNAs when compared to vector control, the RBM3-overexpressing DLD1 cells showed 551 upregulated and 556 down-regulated lncRNAs (Figure 2A). When comparing the lncRNAs between the two cell lines and taking those that are significantly expressed based on an absolute fold-change greater than 1.5 and a p-value less than 0.05, we identified that 67 lncRNAs were commonly dysregulated in both cell lines with RBM3 overexpression (Figure 2B). Next, to detect lncRNAs that interact with RBM3 protein, we also performed RNA-immunoprecipitation coupled sequencing (RNA-IP seq). We observed 368 upregulated and 746 downregulated RBM3-interacting lncRNAs in RBM3-overexpressing HCT116. Similarly, in DLD1 RBM3 overexpressing cells, there were 2882 upregulated and 1467 downregulated RBM3-interacting lncRNAs when compared to vector control (Figure 2C). Overall, in the RNA-IP seq there were 4228 unique lncRNAs in HCT116 and 934 in the DLD1 cell lines, of which 121 were observed in both cell lines (Figure 2D). The heat plot for the gene body coverage of the RNA-seq across the genome also shows differential expression of lncRNA within RBM3 overexpressing and empty vector cells (Figure S2B). We used Gene Ontology (GO) and LncDisease databases to annotate the significantly expressed lncRNAs for their roles in biological functions and diseases. The LncDisease database provided the association of lncRNA to the ascribed functions and diseases. The significance of the association of the differentially expressed lncRNA with the functions and diseases from these databases was calculated using hypergeometric distribution. GO terms enriched in lncRNA seq for RBM3-overexpressing HCT116 cells compared to control were analyzed by REVIGO^30^. The circle size represents the frequency of GO terms, and the color scale represents log10 p-value. In analyzing the data for significantly enriched GO terms, we found that lncRNAs associated with response to wound healing, VEGF production, and blood vessel remodeling were significantly higher in RBM3 overexpressing cells (Figure 2E and 2F). Thus, RBM3 is associated with differential expression of lncRNAs involved in cell movement and angiogenesis.

**Figure 2:**
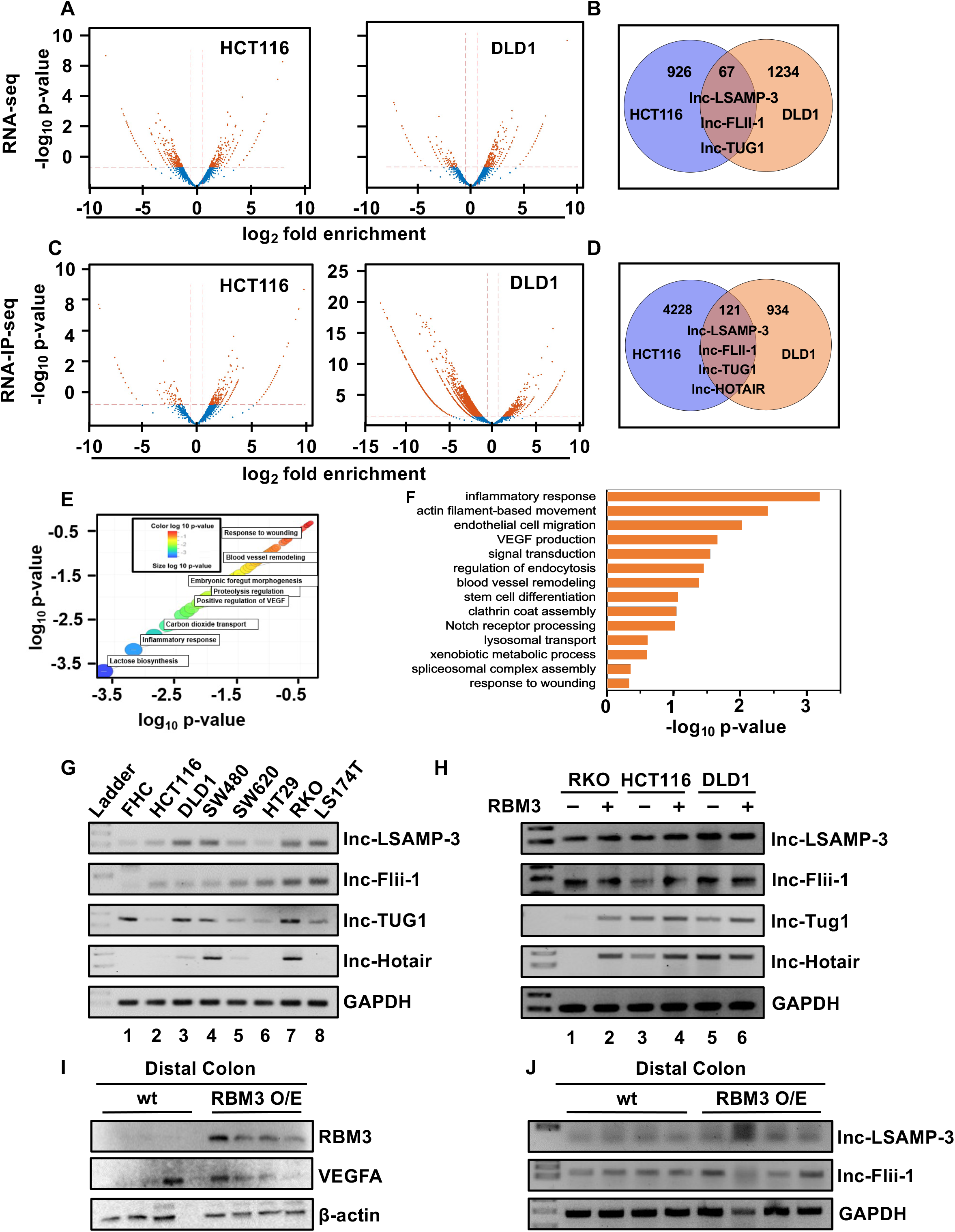
RBM3 causes differential expression of long non-coding RNA. (A) Volcano plots for RNA-sequencing (RNA-seq) showing differentially expressed lncRNA in both HCT116 and DLD1 RBM3 overexpressing (RBM3 O/E) cell lines compared to empty vector (EV) cell lines. (B) Venn diagram depicting the number of co-expressed and uniquely expressed lncRNA found in RNA-seq. (C) The volcano plots for RNA-immunoprecipitation coupled sequencing (RNA-IP seq) showing differentially expressed lncRNA in both HCT116 and DLD1, RBM3 overexpressing cell lines compared to empty vector cell lines. (D) Venn diagram depicting the number of co-expressed and uniquely expressed lncRNA found in RNA-IP seq. (E-F) Plot from REVIGO software showing the Gene Ontology (GO) terms enriched in lncRNA (RNA-seq) for HCT116 RBM3 (E). The graph has been modified to highlight the GO terms and color key. The box plot drawn highlighting enriched GO terms(F). (G) RT-PCR validation of lncRNA identified through RNA-seq in colon cancer cell lines compared to normal FHC cells. (H) RT-PCR validation of lncRNA identified through RNA-seq, in the RBM3 overexpressing (RKO, HCT116, and DLD1) cells, show increased expression of lnc-HOTAIR, lnc-TUG1, lnc-Flii- 1, lnc-LSAMP-3. (I) Western blot analysis for the expression demonstrating levels of RBM3 in the distal colon in representative wild type littermates and RBM3 transgenic mice (RBM3 O/E). (J) RT-PCR analysis demonstrating increased expression of lncRNA lnc-Flii-1 and lnc-LSAMP- 3 compared to GAPDH in representative wild type (WT) littermates and RBM3 overexpressing transgenic mice (RBM3 O/E). Data in Figure 2 are represented as ± SEM. Also, see Supplementary figure 2.

In comparing the common lncRNAs that are upregulated in the RBM3 overexpressing cells and also bind to RBM3, we identified four lncRNAs, of which two are known (lnc-HOTAIR and lnc- TUG1) and two are novel (lnc-Flii-1 and lnc-LSAMP-3) (Figure 2B and 2D). Using the RNAfold software (http://rna.tbi.univie.ac.at/) we determined the predicted secondary structure of the lncRNAs (Figure S2C)^31^. The structure of the lncRNAs is colored by the base-pairing probabilities. Unpaired regions are colored in blue and paired regions are in red Supplementary Figure 2C). We next validated the expression of the four lncRNAs in colon cancer cell lines and a normal colonic epithelial cell line FHC. The expression of lnc-Flii-1, lnc- LSAMP-3 and lnc-HOTAIR were increased in colon cancer cell lines compared to normal FHC cells (Figure 2G). In addition, we observed upregulation of the four lncRNAs in the three cell lines where RBM3 is overexpressed (Figure 2H). Quantitative PCR for lnc-TUG1 revealed a two-fold increase in the lncRNA levels in RBM3 overexpressing cells compared to the empty vector cells (Figure S2D). We then determined if RBM3 expression correlates with lncRNA expression in patient samples. Since gene expression data for lnc-Flii-1 and lnc-LSAMP-3 is not available in the TCGA database, we probed for only lnc-HOTAIR and lnc-TUG1. There was no significant correlation between lnc-HOTAIR and RBM3 levels in interrogating data from TCGA (R=0.076, p=0.054) (Figure S2E). However, a positive correlation was observed between the expression of RBM3 and lnc-TUG1 (r=0.64, p<0.01) (Figure S2F).

To further evaluate the association of RBM3 with lncRNAs, we analyzed colon from transgenic RBM3 overexpressing mice. We generated a RBM3 KnockIn (KI) mice using CRISPR/Cas-mediated genome editing by inserting the “CAG-mouse RBM3 cDNA-Myc-IRES-mCherry-polyA” cassette into intron 1 of *ROSA26* gene locus of C57BL/6 mice in the reverse direction (Figure S2G). The transgenic mice showed an increased expression of RBM3 protein in the distal colon compared to the wild-type litter mates. Also, the RBM3 overexpressing colons had higher levels of VEGFA, a protein that whose expression is regulated by RBM3^12^ (Figure 2I). In addition, the expression of lnc-LSAMP-3 and lnc-Flii-1 was higher in the colon tissue of RBM3 transgenic mice (Figure 2J).

### RBM3 induces angiogenesis, growth, migration and invasion in colon cancer cells

Since VEGF mRNA is increased in response to RBM3 overexpression, we next evaluated the role of RBM3 in angiogenesis by performing the endothelial cell tubule formation assay. HUVEC cells in matrigel were incubated with conditioned media from RBM3-overexpressing HCT116 or DLD1 cells and compared them to empty vector cells. We observed an increase in the tube length in HUVEC cells treated with conditioned media from RBM3-overexpressing HCT116 (p=0.038) and DLD1 (p=0.004) cell lines, indicating that RBM3 facilitates angiogenic tube formation (Figure S3A and 3E). Previously, we reported that RBM3 regulates cancer stemness^19^. To confirm this, we performed spheroid forming assays with the RBM3 overexpressing cells. There was a significant increase in the number of the spheroids with both HCT116 (p=0.047) and DLD1 cells (p=0.001), when RBM3 is overexpressed (Figure S3B, S3C). To determine if RBM3 facilitates cell migration, we subjected spheroids from RBM3 overexpressing or empty vector cells to 2D culture conditions (Figure S3D). We observed that overexpression of RBM3 resulted in increased migration of the cells from 3D spheroids (HCT116 p=0.01, DLD1 p=0.02) (Figure S3E). In addition, RBM3 overexpression increased the migration of cells in the scratch assay, as compared to empty vector in RKO (p=0.001), HCT116 (p=0.023) and DLD1 (p=0.04) (Figure 3F and S3F). Further, RBM3 increased the migration of cells through Boyden chambers and invasion through matrigel in all three cell lines. (Figure 3G,H, S3G,S3H). Thus, RBM3 regulates several hallmarks of cancer including anchorage-independent growth, migration, invasion and inducing angiogenesis.

**Figure 3:**
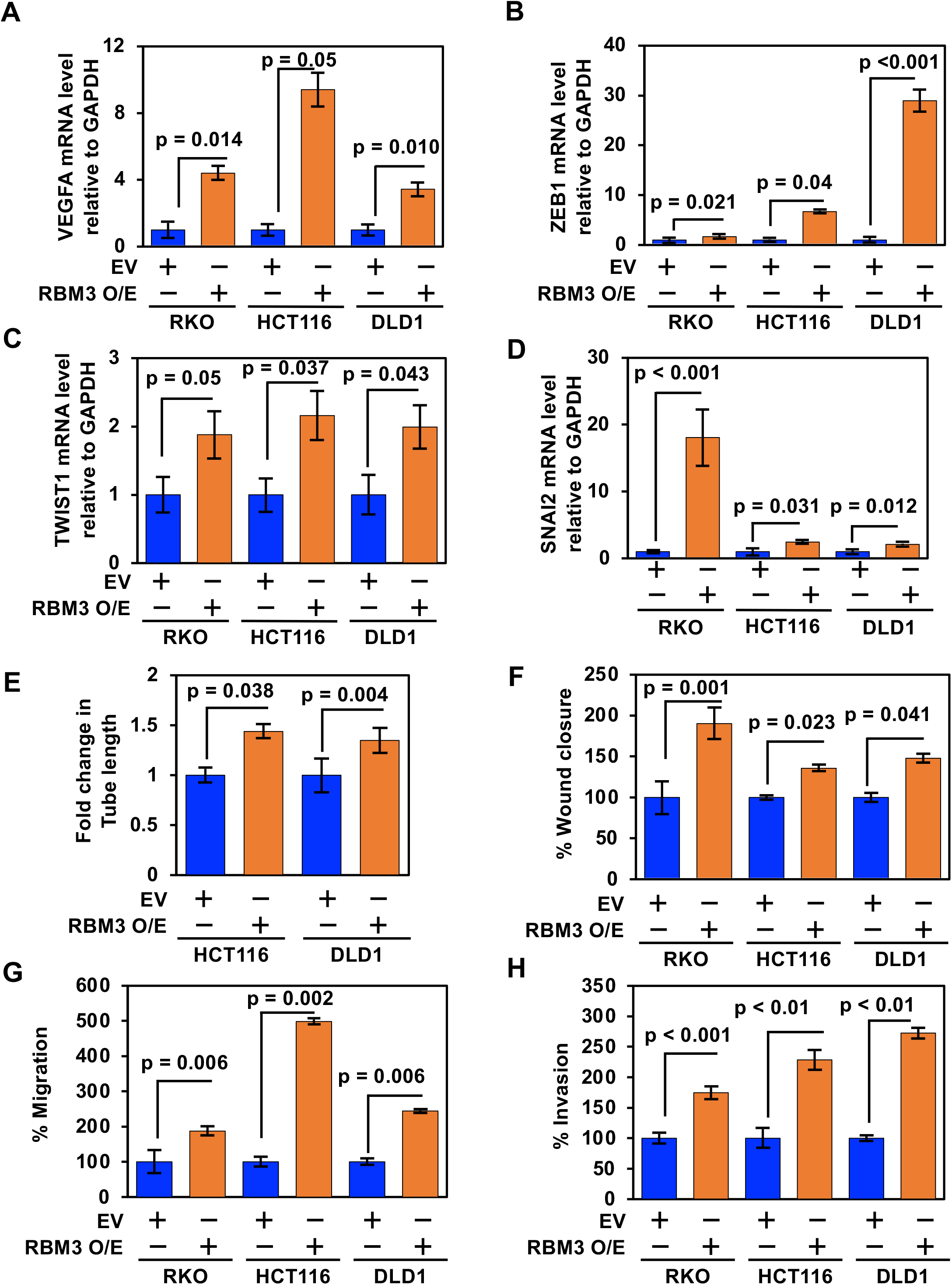
RBM3 induced angiogenesis, migration and invasion. (A-D) Quantitative PCR shows increased mRNA levels of VEGFA, ZEB1, TWIST, and SNAI2 in RBM3 overexpressing cells compared to empty vector cell lines. (E) Plot for tube length of the endothelial tubular network formed by HUVEC cell in (G). Increased tube formation in HUVEC cells treated by condition media from HCT116 (p=0.038) and DLD1 (p=0.004) RBM3 overexpressing compared to empty vector cells (F) Plot showing increase in migration of HCT116 (p=0.001), DLD1 (p=0.02) and RKO (p=0.04) RBM3 overexpressing cells compared to empty vector performed by wound healing assay. (G) Plot showing increase in migration of HCT116 (p=0.006), DLD1 (p=0.002) and RKO (p=0.006) RBM3 overexpressing cells compared to empty vector performed by transwell migration assay. (H) Plot showing increase in invasion of HCT116 (p<0.001), DLD1 (p<0.01) and RKO (p<0.01) RBM3 overexpressing cells compared to empty vector performed by transwell invasion assay. Data in figure 3 are represented as ± SEM. Also, see Supplementary figure 3.

### RBM3 regulated lncRNAs interacts with mRNA involved in angiogenesis and migration

LncRNAs regulate mRNA at various levels including post-transcriptional regulation ^22, 23, 28^. Since, RBM3 induced the expression of VEGFA, ZEB1, SNAI2, and TWIST, we next determined whether these could be putative targets for lnc-HOTAIR, lnc-TUG1, lnc-Flii-1, and lnc-LSAMP-3. For this, we performed *in silico* analyses using the IntaRNA software^32^. For both lnc-Flii-1 and lnc-LSAMP-3, interactions were identified with the 5’- and 3’- untranslated regions of VEGFA, ZEB1, SNAI2, and TWIST mRNA (Table S4A-D, Figure S4A-H). The position of the lncRNA interactions with the four mRNAs along with the minimal energy associated with the interaction is shown. Since lncRNA were predicted to interact with mRNA at the 5’UTR, we determined if there were binding sites for lncRNA on 5’UTR regulatory elements. VEGFA mRNA has several regulatory elements, notable of these are the two Internal Ribosome Entry Sites (IRES) present on the 5’UTR. The two IRES on VEGFA mRNA are the IRES-A that drives the expression of the AUG-initiated forms and IRES-B which drives expression of the CUG-initiated isoforms and helps in cap-independent translation^9^. We determined the predicted structure of IRES-A and IRES-B at the 5’UTR of VEGFA mRNA by RNAfold software. The interactions of the lncRNA on these structures were then highlighted. We observed lnc-Flii-1 and lnc-LSAMP- 3 interaction with the IRES-B (Figure 4A), while lnc-HOTAIR and lnc-TUG1 interactions were with IRES-A (Figure S4I). We also determined whether lncRNA interacts with 5’UTRs of transcripts that undergo cap-dependent translation. For this, we chose ZEB1 mRNA because expression is upregulated in RBM3 overexpressed cells, and the mRNA does not have an IRES. We found that both lnc-Flii-1 and lnc-LSAMP-3 interact with ZEB1 5’UTR (Figure 4B). This suggests that the lnc-RNAs can bind mRNAs that encode either cap-dependent or independent translation.

**Figure 4:**
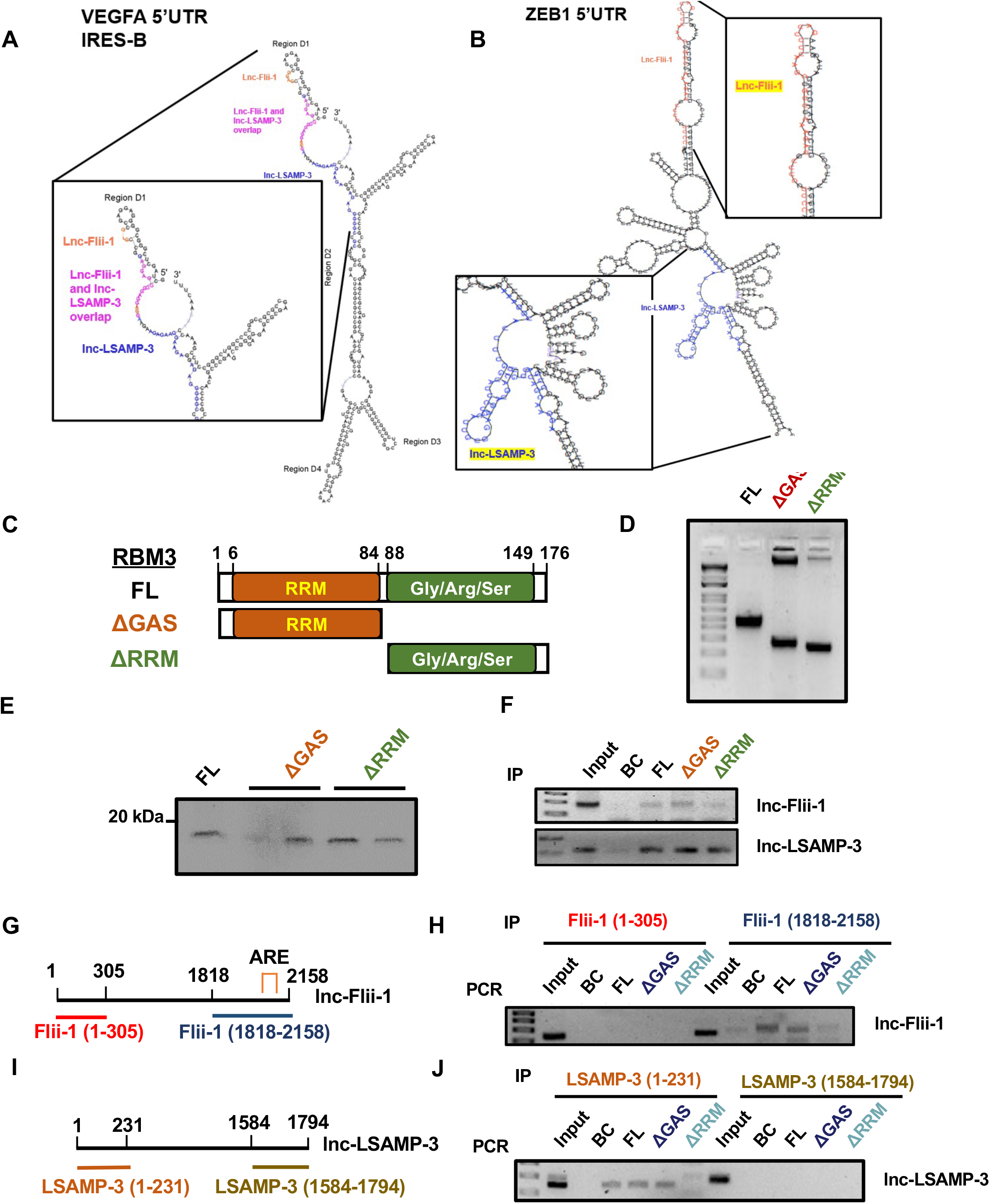
RRM domain of RBM3 associates with lncRNAs. (A-B) Predicted Minimum Free Energy (MFE) secondary structure of VEGFA IRES-B in 5’UTR and ZEB1 5’UTR using the RNAfold web server and modified to highlight the interactions between VEGFA IRES-B and lnc- LSAMP-3, lncFlii-1, and RBM3. (C)Schematic of the RBM3 truncations. *In vitro* protein translation was done for RBM3 full length (FL), GAS domain truncation (ΔGAS), RRM domain truncation (ΔRRM). (D) The PCR products depicting the full length (FL), GAS truncation (ΔGAS), RRM domain truncation (ΔRRM) run on gel. (E) Western blot depicting the full length (FL) and two translation reaction each for GAS truncation (ΔGAS) and RRM truncation (ΔRRM) for RBM3 protein. (F) Immunoprecipitation was done using full length (FL), GAS truncation (ΔGAS), RRM truncation (ΔRRM) protein. PCR for lnc-Flii-1 and lnc-LSAMP-3 was done after pulldown. Whole cell cDNA was used as input and flag beads for bead control (BC). We found that lnc-Flii-1 and lnc-LSAMP-3 were pulled down by full length, RRM and GAS domain for RBM3. (G) Schematic for lnc-Flii-1 truncation showing the invitro transcribed region on 5’ end Flii-1(1-305) and 3’ end Flii-1(1818 to 2158) containing the AU-rich elements. (H) Immunoprecipitation followed by PCR of the lnc-Flii-1 5’ and 3’ region incubated with either FL, ΔGAS and ΔRRM truncation of RBM3 using flag beads. The PCR was performed using primers specific for 5’ end and 3’ end region. The 5’ end of end Flii-1(1-305) was not pulled down by any truncations, while the 3’ end Flii-1(1818 to 2158) was pulled down by FL, ΔRRM and ΔGAS. (I) Schematic for lnc-LSAMP-3 truncation showing the invitro transcribed region on 5’ end LSAMP-3 (1-231) and 3’ end LSAMP-3 (1584 to 1794) containing the AU-rich elements. (J) Immunoprecipitation followed by PCR of the lnc-LSAMP-3 5’ and 3’ region incubated with either FL, ΔGAS and ΔRRM truncation of RBM3 using flag beads. The PCR was performed using primers specific for 5’ end and 3’ end region. The 3’ end of end LSAMP-3 (1584 to 1794) was not pulled down by any truncations, while the 5’ end LSAMP-3 (1-231) was pulled down by FL, ΔRRM and ΔGAS. Data in Figure 4 are represented as ± SEM. Also, see Supplementary figure 4.

### RRM domain of RBM3 interacts with the lnc-LSAMP-3 and lnc-Flii-1

To identify the mechanism of the RBM3-lncRNA interaction, we performed structure-function studies. As RBM3 has two functional domains, an RNA recognition motif (RRM) motif and a C-terminal glycine-arginine-serine rich domain ^14^, we generated truncation mutant constructs that results in expression of only one of these domains (Figure 4A). Primers for the truncation were made to amplify the full length (FL), GAS truncation (ΔGAS), and RRM domain truncation (ΔRRM) with an N-terminal FLAG tag (Figure 4C) and subjected to *in vitro* transcription and translation to generate proteins. After confirming the expression of the full length (FL), GAS truncation (ΔGAS), and RRM truncation (ΔRRM) mutants of RBM3 (Figure 4D), the proteins were incubated with whole cell extracts to identify protein:lncRNA interactions. Following immunoprecipitation for FLAG, we performed RT-PCR for lnc-Flii-1 and lnc-LSAMP-3 with the immunoprecipitate. Both lnc-Flii-1 and lnc-LSAMP-3 binds to the N-terminal RRM domain and the C-terminal GAS domain (Figure 4E). On the other hand, there was no pulldown of lncRNA in bead control (BC) group. RNA binding proteins regulate mRNAs by binding to specific motifs^33^. RBM3 is known to bind to at least one such motif which is the AU-rich element (ARE)^12^. Analyzing the sequence of the lncRNAs, we found AREs at the 5’ end of lnc-LSAMP-3 and at the 3’ end of lnc-Flii-1. Next, we made truncations of the lncRNA removing either 5’ end or the 3’ end. For lnc-Flii-1, we generated truncation constructs containing nucleotides 1-305 and 1818- 2158 (Figure 4G). For lnc-LSAMP-3, we generated truncation that included nucleotides 1-231 and 1584-1794 (Figure 4I). We generated the truncated lncRNA by *in vitro* transcription, and then mixed the RNA with either RBM3 full length protein or the RRM and GAS truncated proteins. Following immunoprecipitation for the protein, RNA interactions were determined by RT-PCR using primer specific to the 5’ end or 3’ end of each lncRNA. We found that the 3’ end of lnc-Flii- 1 interacted with the full length and RRM domain of RBM3, while no interaction was seen in the 5’ end to any RBM3 domain (Figure 4H). For the lnc-LSAMP-3, the 5’ end interacted with RBM3 and the RRM domain, while the 3’ region of lncRNA did not interact with RBM3 (Figure 4J). Since RRM3 bound to ARE sequences in the 5’- and 3’ ends of lnc-LSAMP-3 and lnc-Flii-1 lncRNAs, respectively, these results suggest that the RRM sequence in RBM3 is required for binding to lncRNAs.

### RBM3 overexpression increases tumor growth in the xenograft model

To evaluate the role of RBM3 on tumor growth *in vivo*, we monitored longitudinal growth of tumor xenografts generated with RBM3-overexpressing and vector control, HCT116 and DLD1 cells. We observed that RBM3 overexpression increased the weight and volume of HCT116 (weight p=0.023 and volume p<0.05) and DLD1 (weight p<0.001 and volume p<0.01) xenograft tumors, as compared to empty vector xenografts (n=10 per group, Figure 5A-D). We next validated the expression of lnc-LSAMP-3 and lnc-Flii-1, along with the mRNAs, VEGFA, ZEB1 and TWIST in the tumor xenograft tissues. There were increased levels of both lnc-Flii-1 and lnc-LSAMP-3 in RBM3-overexpressing HCT116 and DLD1 xenografts (Figure 5E). In addition, there were increased levels mRNA transcripts of angiogenesis and EMT markers, VEGFA, ZEB1 and TWIST, when compared to empty vector in the RBM-overexpressing xenografts (Figure 5F). RBM3 overexpressing tumors had higher levels of VEGF protein compared to the empty vector in both HCT116 and DLD1 xenografts (Figure 5G). Immunohistochemistry was performed to assess the protein levels of RBM3 along with markers for proliferation and angiogenesis. RBM3 overexpressing xenografts expressed higher protein levels of PCNA and CD31 compared to the empty vector xenografts (Figure 5H, 5I, S5A and S5B). These data suggest that RBM3 modulates lnc-Flii-1 and lnc-LSAMP-3, facilitating tumor growth, EMT, and angiogenesis.

**Figure 5:**
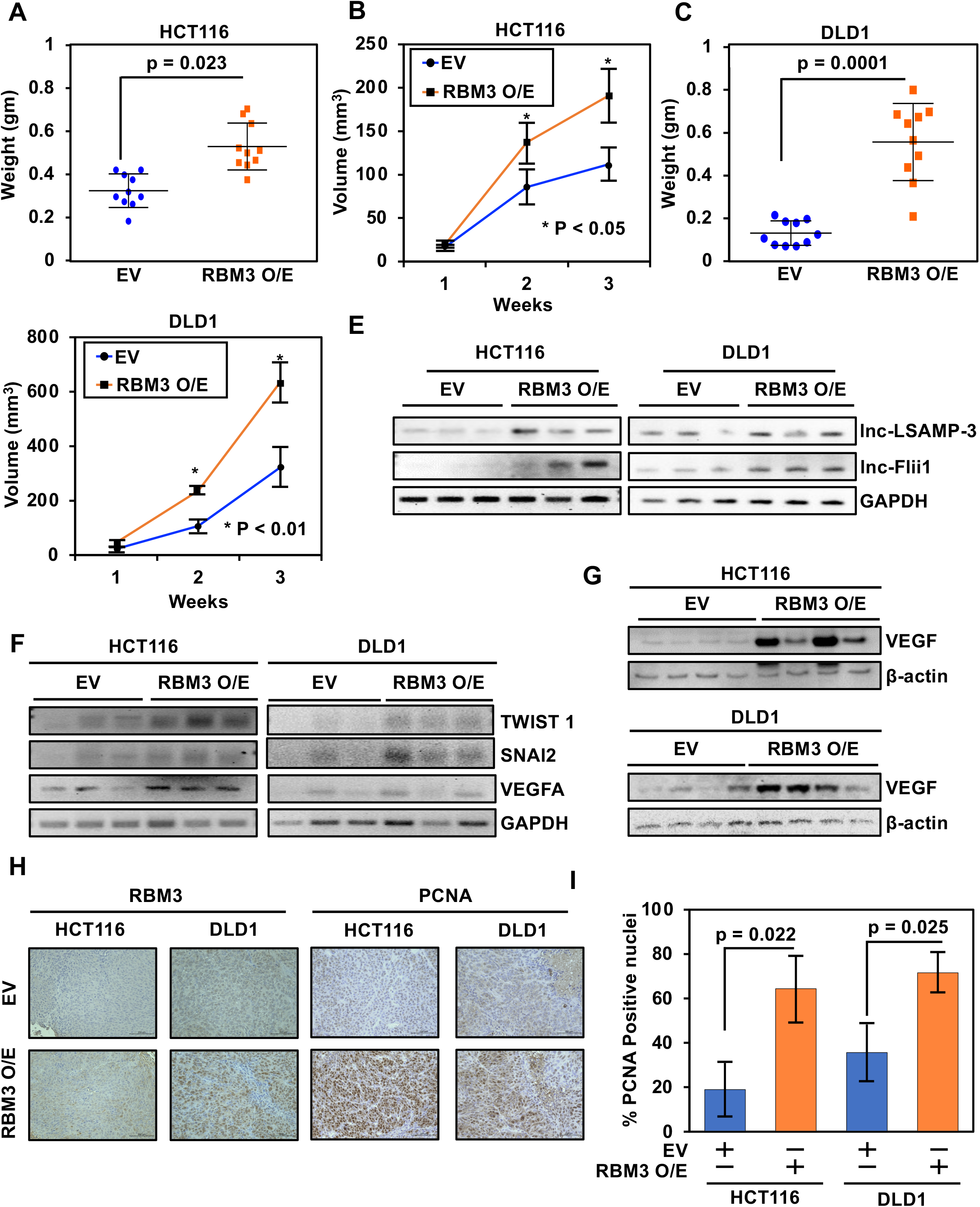
RBM3 overexpression increases tumor growth in the xenograft model. (A-D) The plot of tumor weights (A, C) and tumor volumes (B, D) from HCT116 and DLD1 empty vector and RBM3 overexpressing xenografts. The tumor volume and weight were significantly increased in RBM3 overexpression compared to control (p<0.05) (E) RT-PCR validation of lncRNA in the tumor xenograft tissues, in both HCT116 and DLD1 xenografts. RBM3 overexpressing xenografts show an increase in lnc-Flii-1 and lnc-LSAMP-3 expression compared to empty vector. (F) RT-PCR validation for increased expression of TWIST1, SNA2 and VEGFA in HCT116 and DLD1 empty vector and RBM3 overexpressing xenograft tumors tissues. (G) Western blot analysis of tumors from HCT116 and DLD1 xenograft tissues for VEGF expression. (H) Increassed RBM3 and PCNA levels in HCT116 and DLD1 RBM3 overexpressing xenograft tissues as compared to empty vector as seen by immunohistochemistry analysis. (I) Increase in percentage of PCNA positive nuclei in RBM3 overexpressing HCT116 (p=0.022) and DLD1 (p=0.025) xenograft tissues immunohistochemistry. The data for figure 5 are presented as the means ± SEM, (n=10). Also, see Supplementary figure S5.

### Loss of RBM3 decreases tumor progression *in vitro* and *in vivo*

Having tested the tumor-promoting effects of overexpressing RBM3, we sought to validate these findings by knocking down RBM3 in colon cancer cells. Using shRNA in the HCT116 cells, we achieved over a 50% reduction in RBM3 levels (Figure 6A). There was a corresponding reduction in lnc-LSAMP-3 and lnc-Flii-1 levels, in HCT116 RBM3 knocked down clones compared to the scrambled control cells (Figure 6B). We then assessed the proliferative capacity of the RBM3 shRNA knockdown cell clones over the scramble control cells. RBM3 knockdown significantly reduced the proliferation rate of the three shRNA cell clones (p<0.05) when compared to the scramble controls (Figure 6C). We also assessed the effect of RBM3 knockdown on the colony forming capacity. The shRNA knockdown clones of RBM3 showed a decrease in the number and size of colonies as compared to the scramble control (Figure S6A- S6C). RBM3 knockdown also significantly reduced the migration and invasion of cells (shRNA- 2 migration p=0.04 and invasion p<0.001, and shRNA-5 migration p=0.04 and invasion p=0.001) (Figure 6D, 6E, S6D, S6E). Knockdown of RBM3 also significantly impaired the increase in tumor xenograft weight (p=0.007) and volume (p<0.05) compared to that of scrambled control tumors (Figure 6F and 6G). RBM3 knockdown also attenuated expression of lnc-LSAMP-3 and lnc-Flii-1 in the xenograft tumors (Figure S6E). These data, taken together, demonstrate that RBM3 regulates tumor proliferation, migration and invasion. Further, RBM3 may facilitate tumor growth through regulation of lnc-LSAMP-3 and lnc-Flii-1.

**Figure 6:**
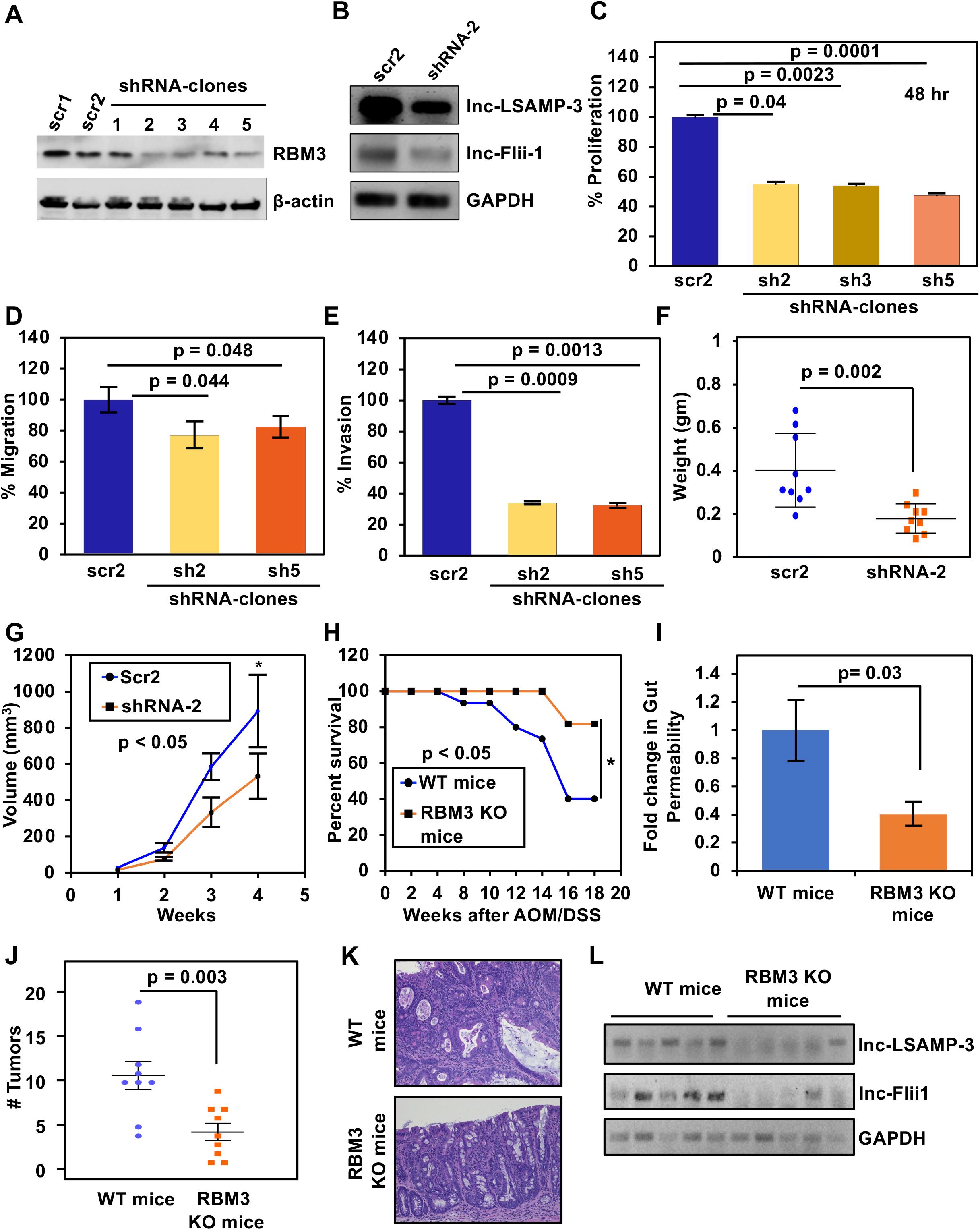
RBM3 knockdown decreases lncRNA expression causing decreased cell migration, invasion and tumor growth in the xenograft model. (A) Protein expression of RBM3 decreases in shRNA knockdown clones (2,3,5) compared to scramble (scr1,scr2) in the HCT116 cell line by Western blot analysis. (B) RT-PCR validation for the decreased expression of lnc-LSAMP-3 and lnc-Flii-1 in HCT116 RBM3 shRNA knocked down clone compared to scramble. (C) Plot for decrease in percentage proliferation of the RBM3 shRNA knockdown clones over empty vector for sh2 (proliferation= 55.06%, p=0.04), sh3 (proliferation= 53.8%, p=0.0023), sh5 (proliferation= 47.45%, p=0.0001). (D) The plot showing decrease in percent migration in HCT116 shRNA knocked down clones (sh2 p=0.044 and sh5 p=0.048) compared to scramble performed by Transwell migration. (E) The plot showing decrease in percent invasion in HCT116 shRNA knocked down clones (sh2 p=<0.001 and sh5 p=0.001) compared to scramble performed by Transwell invasion assay. (F and G) The plot of tumor weights (F) and tumor volumes (G) from HCT116 scramble (scr2) and RBM3 shRNA knockdown (shRNA-2) xenografts. Both tumor volume and weight were significantly reduced in RBM3 knockdown compared to scramble control (p<0.05). (H) Kaplan-Meier analysis showing increased percent survival after the AOM/DSS induced carcinogenesis in cre inducible RBM3 knockout (KO) compared to wild type (WT) mice. (I) Plot showing decreased intestinal permeability in RBM3 knockout mice as compared to the wild type littermates (p=0.04). (J) The plot showing decreased number of tumors in the distal colon observed after the AOM/DSS induced carcinogenesis in cre inducible RBM3 knockout compared to wild type mice (p=0.007). (K) Representative image of H and E staining of the distal colon with tumors after the AOM/DSS induced carcinogenesis in RBM3 knockout and wild type mice. (L) Decrease in expression of lnc-LSAMP-3 and lnc-Flii-1 seen by PCR analysis in RBM3 knockout mice as compared to the wild type mice. The data for figure 6 are presented as the means ± SEM, (mice n=10). Also, see Supplementary figure 6.

Given the reduction in xenograft growth following RBM3 knockdown, we next determined the effect of loss of endogenous RBM3 on colon tumorigenesis. For this, we generated a mouse where exons 2-6 were flanked by loxP sequence (Figure S6F). After confirmation of the loxP insertion by PCR, we crossed the mice with mice carrying the colon-specific promoter, CDX2- driving Cre/ER^T^^2^ gene^34^ (Figure S6F and S6G). Subsequently, we administered tamoxifen, resulting in knockout of RBM3 specifically in intestinal epithelial cells (Figure S6H). To determine the effect of RBM3 loss on tumorigenesis, we treated the mice with azoxymethane and dextran sodium sulfate (AOM/DSS). AOM is a carcinogen frequently used to induce colon cancer in mice^35^. In this model, mice develop colitis which then rapidly progresses to cancer in the colon^35^. Wild-type and RBM3 knockout mice (n=10, per group) were treated with a single AOM intraperitoneal injection, followed by three rounds of oral DSS (Figure S6I). Mice were monitored for external signs of rectal prolapse as an indicator of colitis. In as early as 14 weeks post carcinogen induction, 86.6% of the wild-type mice developed rectal prolapse while only 36.3% of RBM3 knockout mice developed the condition. At 20 weeks post-treatment, only 40% of wild-type mice, compared to 80% of RBM3 knockout mice survived the AOM/DSS treatment (Figure 7H). Perturbation in the intestinal mucosal barrier is associated with intestinal diseases especially in inflammatory bowel disease (IBD). A breach in the intestinal barrier can exacerbate inflammation and promote tumor progression^36^. Therefore, we investigated the intestinal permeability of WT and RBM3 knockout mice. There was an over two-fold reduction in intestinal permeability in the RBM3 knockout mice (Figure 7I). Thus, RBM3 plays an important role in the progression of colitis to cancer, severely impacting survival.

**Figure 7:**
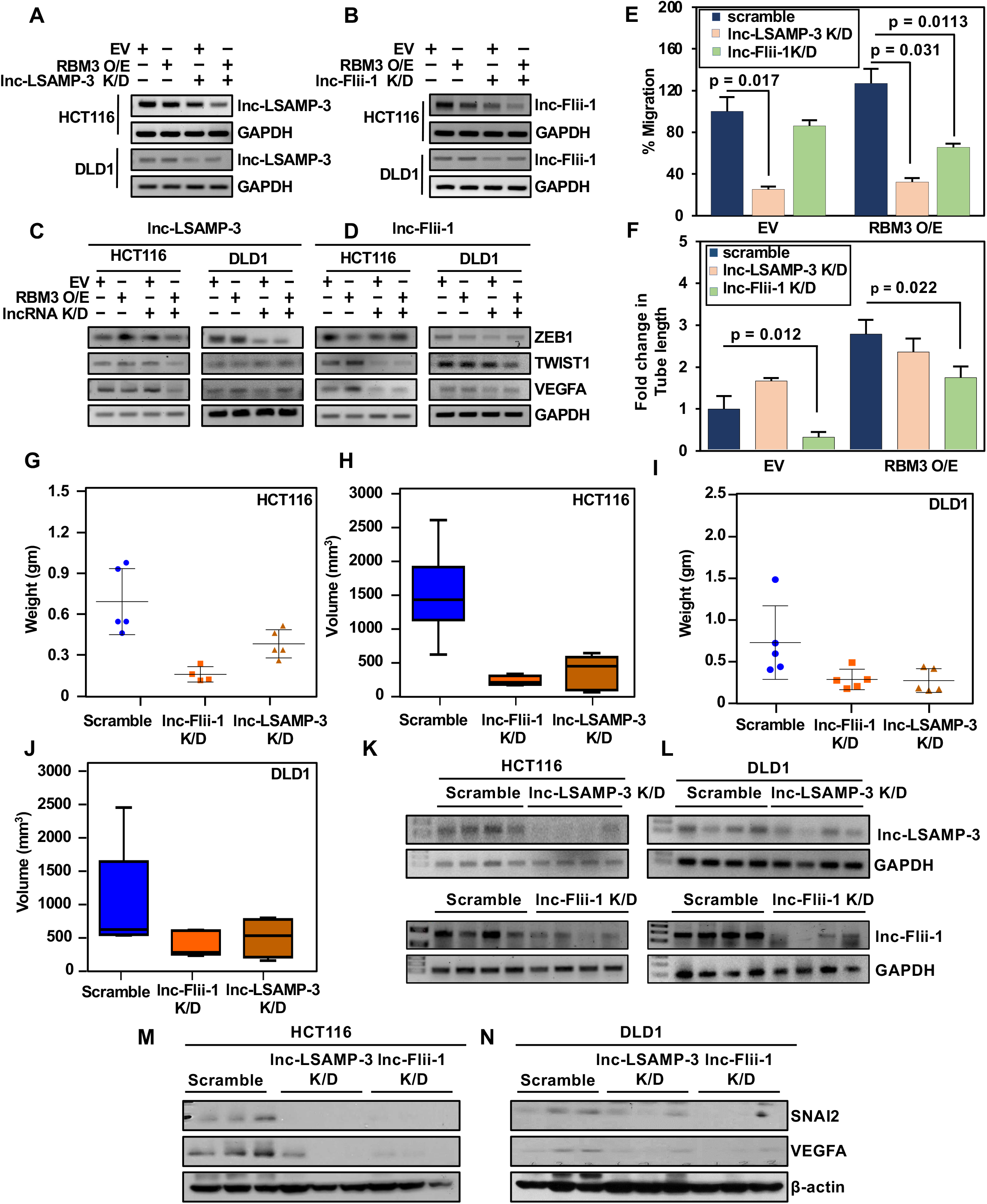
RBM3 induces cancer cell migration and Invasion through lncRNA. (A and B) RT-PCR validation of lnc-LSAMP-3 and lnc-Flii-1 knocked down in HCT116 (A) and DLD1 (B) RBM3 O/E and empty vector cells. (C and D) RT-PCR validation of VEGFA, ZEB1 and TWIST in HCT116 (C) and DLD1 (D) RBM3 O/E and empty vector cells, after lncRNA (lnc-Flii-1 and lnc-LSAMP-3) knockdown. (E) Plot showing decreased percent migration of HCT116 empty vector and RBM3 overexpressing cells having knockdown of lncRNA lnc-LSAMP-3 (Empty vector p=0.017, RBM3 O/E p=0.031) and lnc-Flii-1(Empty vector p=ns, RBM3 O/E p=0.011) performed by scratch plate assay. (F) Plot showing decreased tube length of the endothelial tubular network formed by HUVEC cell treated by condition media from knockdown of lncRNA lnc-LSAMP-3 and lnc-Flii-1(Empty vector p=0.012, RBM3 O/E p=0.022) in HCT116 cells. (G-J)The plot showing decrease in tumor weight (G, I) and tumor volumes (H, J) from HCT116 and DLD1 xenografts treated with intratumoral injection of si+ LNA gapmer for lnc-Flii-1 knockdown and lnc-LSAMP-3 knockdown compared to control (p< 0.05). (K and L) RT-PCR validation for the decreased expression of lncRNA (lnc-LSAMP-3 and lnc- Flii-1) in the tumor xenograft of HCT116 (K) and DLD1(L) treated with intratumoral injection of si+ LNA gapmer for lnc-Flii-1 knockdown and lnc-LSAMP-3 knockdown compared to control. (M and N) Western blot showing decreased protein expression of SNAI2 and VEGF in HCT116 (M) and DLD1 (N) tumor xenograft treated with intratumoral injection of si+ LNA gapmer for lnc- Flii-1 knockdown and lnc-LSAMP-3 knockdown compared to control. The data for figure 7 are presented as the means ± SEM. Also, see figure Supplementary figure

Gross examination of the distal colon from mice demonstrated a significant attenuation in the number of tumors in the colon of RBM3 knockout mice compared to that from wild-type animals. Further, 100% of the wild-type mice presented with tumors in the colon in contrast with 80% in RBM3 knockout mice. The average number of tumors per wild-type mouse (10.77±1.26), was about three times higher than in RBM3 knockout mice (3.2±0.85) (Figure 7J and S6J). The histology of the distal colon from both WT and RBM3 knockout mice treated with AOM/DSS was analyzed. Hematoxylin and eosin staining also showed pathological features indicative of injury and inflammation (Figure S6J). There were two RBM3 knockout mice that had no tumors and the histology showed normal mucosa, while one mouse had low-grade adenocarcinoma. The remaining RBM3 knockout mice showed tumors that were histopathologically indistinguishable from those of WT mice. These tumors were composed of intratumoral crypt abscess and glands consistent with a well-differentiated adenocarcinoma (Figure 7K and S6K). However, the number of tumor foci and the average size of foci was decreased in RBM3 knockout mice compared to those in WT mice (Figure S6L). The expression of lnc-LSAMP-3 and lnc-Flii-1 also decreased in the tumors of RBM3 KO mice compared to the WT tumors (Figure 7L). These cumulative findings demonstrate that RBM3 regulates colon cancer progression, and that lnc-LSAMP-3 and lnc-Flii-1 are involved in the process.

### lncRNA knockdown decreases *in vitro* cell migration, angiogenesis and tumor growth in the xenograft model

To directly address the role of lncRNAs in RBM3-induced cell migration and angiogenesis, we used a combination of siRNA and locked nucleic acid oligomers to knockdown lnc-Flii-1 and lnc- LSAMP-3. The lncRNAs were knocked down in the RBM3 overexpressing HCT116 and DLD1 cells, and empty vector cells. The extent of lncRNA knockdown was confirmed by PCR. (Figure 7A and 8B). Knockdown of either lnc-Flii-1 and lnc-LSAMP-3 also resulted in reduced expression of VEGFA, ZEB1 and TWIST in both HCT116 and DLD1 cells despite RBM3 overexpression (Figure 7C and 7D). Furthermore, knockdown of lnc-LSAMP-3, significantly reduced migration in both empty vector and RBM3 overexpressing cells, while lnc-Flii-1 knockdown had a significant reduction in migration only in the RBM3 overexpressing cells (p=0.0113) (Figure 7E). Interestingly, RBM3-induced angiogenesis was effectively mitigated by lnc-Flii-1 knockdown (p=0.012) but not lnc-LSAMP-3 knockdown (Figure 7F). These data suggest that RBM3 exerts its tumor-promoting effects through differential use of lncRNA.

We next tested the role of lnc-LSAMP-3 and lnc-Flii-1 in tumor growth *in vivo*. HCT116 and DLD1 tumor xenograft-bearing mice were treated intratumorally with a combination of siRNA and LNA gapmer targeting lnc-LSAMP-3 or lnc-Flii-1 (Figure S7A). Attenuation of both lnc- LSAMP-3 and lnc-Flii-1 reduced tumor volume and weight as compared to tumors treated with scramble controls in HCT116 and DLD1 xenografts (Figure 7G-J). The tumors treated with siRNA+LNA gapmer showed decreased expression of lnc-Flii-1 and lnc-LSAMP-3 (Figure 7K and 7L). The tumors also showed a corresponding decrease in the protein expression of VEGF and Slug (SNAI2) (Figure 7M and 7N). Thus, lnc-LSAMP-3 and lnc-Flii-1 regulate colon cancer growth in part through the modulation of angiogenic factor VEGF, and EMT marker Slug.

## Discussion

Deregulation of RBPs have been shown to affect the various hallmarks of cancer leading to tumorigenesis^1, 37^. Previously, we have demonstrated that increased expression of RBM3 resulted in upregulation in cell proliferation, while loss of RBM3 resulted in cells undergoing mitotic catastrophe^12^. Now, we have investigated its role in stemness and the EMT phenotype. While RBM3 overexpression increased stemness, cell movement, and angiogenesis, this was reduced by shRNA knockdown of the protein. Also, while RBM3 overexpression increased xenograft tumor, RBM3 knockdown decreased the growth. EMT markers and VEGFA were increased with RBM3 expression contributing to increased tumor growth. ZEB1, Snail, Slug, and TWIST transcription factors orchestrate EMT by repressing adhesion protein E-cadherin and increasing mesenchymal phenotypes^38^. Several studies have shown that EMT may have a role in switching dormant cells to a proliferative phenotype^39, 40^. In addition, transcription factors ZEB1 and TWIST have previously been shown to promote CSC traits in cancer^41, 42^. It is well established that EMT and stemness contribute to tumor progression and metastasis. RBM3 can increase the stem-like phenotype in colon cancer cells, and we demonstrated this by increased spheroids and side population compared to parental controls. Further, we demonstrate an increased expression of the stem cell markers including LGR5, DCLK1 and an increase in the CD44^Hi^/CD24^Low^ population ^19^.

Although mRNA is the primary substrate for RBPs, recent studies have shown that RBPs also interact with non-coding RNAs^43^. For instance, RBP, hRNP-K interacts with several lncRNAs to affect development and oncogenesis^44^. In addition, MATR3 interacts with lncRNA SNHG1 to increase progression in neuroblastoma^45^. LncRNAs can either act as oncogenes or as tumor suppressors. Several lncRNAs including HOTAIR and TUG1, have been implicated in cancer^46–48^. Moreover, lncRNAs including HOTAIR and others have the ability to regulate EMT in CSCs, contributing to drug resistance, tumor progression, relapse and metastasis^49, 50^.

Previously lncRNAs were shown to be involved in colon cancer proliferation, invasion, and metastasis^51^. In this study, we utilized RNA-immunoprecipitation coupled sequencing to identify non-coding RNAs associated with RBM3. We focused on lncRNAs because it is an understudied area. In our study, we found two novel lncRNAs Flii-1 and LSAMP-3 along with HOTAIR and TUG1 to be upregulated in RBM3 overexpressing cells. HOTAIR and TUG1 have previously been shown to be involved in stemness and EMT. In breast cancer HOTAIR was observed to be high in CD44^+^CD24^-^ stem cells when compared to non-stem cells^52^. Similarly, HOTAIR was found to be upregulated in stem cell population of hepatocellular and lung^53^. HOTAIR has been shown to regulate Snail, Slug, E-cadherin, and MMPs to promote EMT^49, 54^. TUG1 has also been shown to affect EMT by regulating ZEB2^54, 55^. We describe here for the first time that lncRNAs, Flii-1 and LSAMP-3 also regulate ZEB1, Twist1 and Slug mRNAs. In addition, these lncRNAs facilitate angiogenesis by affecting VEGFA expression. The lncRNAs interact with *cis*-acting structural elements in the 5’UTR of the mRNAs of both IRES encoding or non IRES mRNAs. Our in-silico analysis shows that HOTAIR and TUG1 interact with the IRES-B, while Flii-1 and LSAMP-3 interact with IRES-A in the 5’UTR of VEGFA mRNA. Our data suggest that these lncRNAs regulate mRNA through both cap-dependent and independent mechanisms.

The role of RBM3 in tumorigenesis has not been studied in animal models, cre-induible knockout mouse model and tumor xenografts. While knockout of RBM3 resulted in reduced tumor formation in the AOM/DSS model, there was also a reduction in tumor xenografts when RBM3 levels were suppressed using specific shRNA. These data suggest that RBM3 may have an important role in the development of colitis induced cancer. Previously, Matsuda et.al generated mice deficient in RBM3 to study the role of RBM3 in immunity^56^. However, they found no abnormalities in the RBM3 knockout mice, although mouse embryonic fibroblasts (MEFs) generated from these mice showed decreased proliferation compared to control MEFs. More importantly, they found that RBM3 caused G2-phase arrest in the MEFs, similar to what we determined in colon cancer cells^12, 56^. To explore the functional interplay between RBM3 and lncRNAs, we performed knocked down studies. Despite RBM3 overexpression, knockdown of either lncRNA LSAMP-3 or Flii-1 resulted in reduced cellular movement. Further, conditioned media from lncRNA modulated colon cancer cells induced endothelial tube formation to a lesser extent than conditioned media from control cells. In addition, knockdown of the lncRNAs resulted in reduced tumor xenograft growth, coupled with decreased expression of VEGF and Slug protein, suggesting that the lncRNAs modulate EMT and angiogenesis. Furthermore, these data suggest that RBM3 induced EMT and angiogenesis occurs in part through the lnc-LSAMP-3 and lnc-Flii-1. To determine the effect of RBM3 on endogenous expression of these lncRNAs, we employed both the RBM3 KnockIn transgenic mice and the cre-specific knockout mice. While we were able to show that HOTAIR and TUG1 are upregulated in the RBM3 overexpression but reduced in the knockouts, we were unable to perform such studies with LSAMP-3 and Flii-1 because a lack of identified orthologs. However, in scanning the mouse genome, we did identify genomic sequences with similarity to these lncRNAs. RT-PCR analyses demonstrates the presence of these transcripts in mouse colonic tissues, which was significantly reduced in the RBM3 knockout mice even following treatment with AOM/DSS, while it is upregulated in the RBM3 transgenic mice. Further studies involving knockout of these two sequences in the mouse will be required for unequivocally determining if they are mouse orthologs of human LSAMP-3 and FLii-1.

Deletion analyses suggests that the RRM domain in RBM3, interacts with LSAMP-3 and Fliii-1. In fact, RBM3 bound a region that encodes AU-rich sequences in the two lncRNAs. Further mutagenesis studies are required to demonstrate that the RRM in RBM3 interacts with the AU-rich sequences in the lncRNAs. Nevertheless, this is the first demonstration that RRM-containing RNA binding proteins modulate lncRNA function through interaction with AU-rich sequences. An interesting observation is that the GAS domain in RBM3 also interacts with LSAMP-3 RNA, suggesting that this domain also has RNA binding properties. Again, further studies are required to delineate the differences in RNA binding between the RRM and GAS domains in RBM3, and the role of this binding on tumorigenesis. Interestingly, RBM3 encodes only one RRM domain. Previous studies with ELAV family proteins have suggested that a combination of two RRMs has a higher affinity of binding to AU rich elements^57, 58^. The possibility therefore exists that the GAS domain in RBM3 enhances RRM binding function, and this can be further determined by performing additional structure-function studies. Otherwise, RBM3 could interact with other RNA binding proteins and together modulate lncRNA function. In this regard, it should be noted that RBM3 was identified as a HUR interacting partner in colon cancer^12^,. It would be interesting to determine whether RBM3 requires HuR to coordinate with the lncRNAs to modulate stability and translation of VEGFA and EMT encoding mRNAs. Moreover, the current study also raises the question of whether lncRNA function(s) are modulated by coordinated activities of various RNA binding proteins, and if so then understanding the RNA binding protein-lncRNA interaction network that functions in a coordinated fashion to modulate cellular functions would be of high significance.

The role of lncRNA in translational control is not well understood. Recent studies have shown that lncRNAs are present and associated with the ribosomes and poly-ribosomal fractions^27, 59^. Given that RBM3 can also enhance global protein translation by interacting with the 60S ribosomal subunit and increasing the formation of active polysomes^29^, the lncRNA association opens the possibility of additional mechanisms of regulating mRNA translation, which needs further investigation ^25, 26^. The role of RBM3 and associated lncRNA in cancer progression can be elucidated to the translational control of the mRNAs involved in EMT and stemness. In addition, RBM3 overexpression enhances phosphorylation of translation initiation factors eIF4E and 4EBP1. Since 4EBP1 is a translation inhibitor that interacts with eIF4E, and that phosphorylated 4EBP1 loses this ability to interact, thereby releasing eIF4E to enhance mRNA translation^60^, the possibility further exists that there is coo-ordinated activities of all these factors controlled by RBM3 in regulating cap-dependent translation. This may also be the case for cap-independent, IRES-based translation since RBM3 enlists the help of lncRNAs to bind to the IRES and the 3’UTR of VEGF mRNA. Further analysis is necessary to determine whether this is unique to VEGFA or whether other RNAs also undergo this same mechanism of regulation controlled by RBM3 or another RNA binding protein. This notwithstanding, this is the first study that has characterized the coordinated actions of a RNA binding protein and lncRNAs in helping circularization of mRNA, a critical mechanism for the recruitment of ribosomes to facilitate multiple rounds of mRNA translation ^61^. The interaction of RBM3 and lncRNA to mRNA defines the novel way through which ncRNAs could affect the translation. Further studies are needed to evaluate the mechanism and working of this model. Collectively, our data suggest that the association of RBM3 with lncRNA facilitates kissing loop interactions between lncRNA and the mRNA, increasing translation of stemness and EMT related mRNAs driving tumor progression.

## Methods

### Cells and reagents

Human colon cancer cells HCT116, DLD1, RKO and endothelial cell line HUVEC (all cell lines obtained from American Type Culture Collection, at passage 4). The cell lines were grown in DMEM with 4.5 g/L glucose, L-glutamine and sodium pyruvate (Corning, Tewksbury, MA) containing 10% heat-inactivated fetal bovine serum (Sigma-Aldrich, St. Louis, MO) and 1% antibiotic-antimycotic solution (Corning, Tewksbury, MA), at 37°C in a humidified atmosphere of 5% CO2. Following transfection with empty vector and tRBM3 plasmids, cells were cultured as mentioned above with an additional 1mg/mL G-418 (Santa Cruz Biotechnology) and 1µg/mL puromycin (Life Technologies). All the cell lines used in this study were within 20 passages after receipt or resuscitation (∼3 months of non-continuous culturing).

### Animals

Mice used for the xenograft experiments were five-week-old male athymic Foxn1nu mice, purchased from Charles River Laboratory. The mice were maintained with water and standard mouse chow *ad libidum* and used in protocols approved by the Animal Care and Use Committee.

### Colon cancer xenograft model

Athymic Foxn1nu mice were injected with 1 × 10^6^ cells subcutaneously in the flank. Either empty vector or RBM3 overexpressing cells (DLD1 or HCT1), or HCT116 cells stably expressing scramble or RBM3 shRNA were used. The tumors were allowed to grow for three weeks. Tumor volumes were measured with vernier calipers weekly and the volume calculated [(length × width^2^) × 0.5]. At the end of three weeks, the animals were euthanized, and the tumors were weighed and photographed. For lncRNA knockdown, siRNA+LNA gapmer were incorporated into DOPC (1,2-Dioleoyl-*sn*-Glycero-3-Phosphocholine) ^62^. Xenograft tumors were generated by injecting HCT116 and DLD1 cells (1 × 10^6^ cells) subcutaneously into the flanks of male athymic Foxn1nu mice and housed under specific pathogen-free conditions. When tumors reached 150 mm^3^, they were injected intratumorally with 25ul (10 μM) siRNA+LNA gapmer on every third day from day 15 for a total of 5 doses. Tumor volume were measured every third day. We included five mice per group and the experiment was repeated thrice. All tumors were excised after four weeks of treatment and photographed; for each tumor, about half of the tumor was frozen in liquid nitrogen and stored at −80°C, and the rest part was fixed with 10% formalin and embedded in paraffin. The tissue protein and RNA were extracted using RIPA buffer and TRIzol method respectively. The paraffin-embedded tissues were used for immunohistochemical analysis. Tumor volume and weight was plotted and represented as mean±S.E.M.

### RBM3 overexpressing transgenic mice model

RBM3 overexpressing transgenic mice were generated by RBM3 CDS KnockIn (KI) at ROSA26 locus of in C57BL/6 mice. CRISPR/Cas-mediated genome engineering was used to generate the mice from Cyagen Biosciences INC. For the KI model, the “CAG-mouse Rbm3 cDNA-Myc-IRES-mCherry-polyA” cassette was cloned into intron 1 of ROSA26 in reverse direction. To engineer the donor vector, homology arms were generated by PCR using a BAC clone from the C57BL/6J library as a template. Cas9 and gRNA were co-injected into fertilized eggs with donor vector for KI mice production. The pups born were then genotyped by PCR followed by sequencing of PCR product. For experiment littermates or age-matched C57BL/6 mice were used. Mice were genotyped before experiments.

### RBM3 knockout mice model

The RBM3 Cre inducible knockout mice were created from Cyagen Biosciences INC. The Rbm3 gene (NCBI Reference Sequence: NM_016809.6; Ensembl: ENSMUSG00000031167) is located on mouse chromosome X and has seven exons. The ATG start codon is present in exon 2 and the TGA stop codon in exon 6 (Transcript: RBM3-005 ENSMUST00000040010). Exons 2∼6 were selected as the conditional knockout region. To engineer the targeting vector, homology arms and cKO region was generated by PCR using BAC clone RP23-224M4 and RP23-77B24 from the C57BL/6J library as a template. C57BL/6 ES cells were used for gene targeting. The constitutive KO allele was obtained with Cre-mediated recombination after crossing the RBM3 KO mice with CDX2-Cre/ERT2. Mice were genotyped for both RBM3 floxed and CDX2-Cre/ERT2 before experiments. Tamoxifen was used to induce knockout, which was confirmed by PCR. For AOM/DSS experiment littermates or age-matched C57BL/6 mice were used.

### AOM/DSS model in RBM3 knockout transgenic mice

Six to 8-week-old RBM3 KO and age-matched wild type C57BL/6 were used for experimental and control groups. The weight of the mice was recorded on Day 0 and each mouse was injected intraperitoneally (IP) with 10 mg/kg of AOM working solution (1 mg/ml in isotonic saline, diluted from 10 mg/ml stock solution in dH2O kept at -20°C). On day 7, DSS (2.5%) solution was supplied to mice as their drinking water for 5 days (Approximately 250 ml/cage). DSS solution should be replaced in clean bottles three times (every 2-3 days) during this period. On day 14, the cages were switched back to standard drinking water for two weeks. This cycle was repeated on days 28 and 49. A DSS “cycle” consists of one week of DSS in the drinking water followed by 2 weeks of regular (autoclaved) water. The mice were euthanized after 24 weeks from the start of the study. AOM/DSS can cause other disease symptoms like blood in feces, diarrhea, rectal prolapse and dehydration ^63^. Animals were monitored for these symptoms and euthanized when disease symptoms were severe. DSS-induces colitis damages the distal colon; therefore, the entire colon was assessed for tumor burden. The gross tumor burden was recorded and tumors were photographed and excised, about half of the tumor was frozen in liquid nitrogen and stored at −80**°**C, and the rest part was fixed with 10% formalin and embedded in paraffin. The tissue protein and RNA were extracted using RIPA buffer and TRIzol method respectively. The paraffin-embedded tissues were used for immunohistochemistry analysis. We also assayed the gut permeability after the AOM/DSS treatment at the end of the study.

### The Cancer Genome Atlas Data Analysis

TCGA colon adenocarcinoma (COAD) cohort gene expression RNAseq data downloaded using UALCAN Browser ^64^ (http://ualcan.path.uab.edu/ ). The UALCAN browser uses TCGA level 3 RNA-seq and clinical data from 31 cancer types. The UALCAN browser can analyze the relative expression of any gene across tumor and normal samples. It also analyses the expression of the gene in various tumor sub-groups based on individual cancer stages, tumor grade, race, body weight or other clinicopathologic features. Expression levels of RBM3 were designated as transcripts per million in colon adenocarcinoma compared to normal. Boxplot showing relative expression of RBM3 in normal and Colon adenocarcinoma along with statistical significance was generated from ULCAN browser.

### Real-time reverse-transcription PCR analysis

Colon cancer cDNA panel with matched adjacent tissue controls was obtained from Origene (Rockville, MD). Total RNA from cell lines was isolated using TRIzol reagent (Invitrogen, Carlsbad, CA) following manufacturer’s instructions. Two μg RNA was used to synthesize complementary DNA using Superscript II reverse transcriptase and random hexanucleotide primers (Invitrogen, Carlsbad, CA). Individual gene expression was quantified by PCR analysis with complementary DNAs by using a premix containing *Taq* DNA Polymerase (Takara, Shiga Prefecture, Japan). The PCR products were resolved on 2% agarose gel and imaged on BioRad ChemiDoc--XRS+ instrument and analyzed by ImageJ software. The real-time PCR analysis was also performed with the Complementary DNAs using JumpStart Taq DNA polymerase (Sigma-Aldrich, St. Louis, MO) and SYBR green nucleic acid stain (Molecular Probes, Eugene, OR). Crossing threshold values for individual genes were normalized to GAPDH as an internal standard. Changes in mRNA expression were expressed as fold change relative to control. Primers used for the PCR include RBM3, VEGFA, ZEB1, Snai2, TWIST, lnc-HOTAIR, lnc-TUG1, lnc-Flii-1, lnc-LSAMP-3.

### Western blot analysis

For western blot analysis, colon cancer cells were washed with PBS (3 times) and lysed in protein lysis buffer (ThermoFisher Waltham, MA) containing protease inhibitor (Roche, Basel, Switzerland). The resultant lysates were centrifuged at 6000 rpm for 10 mins. Protein (30– 75 µg) was loaded into gels. Cell lysates were subjected to polyacrylamide gel electrophoresis and transferred onto Immobilion polyvinylidene difluoride membranes (Millipore, Bedford, MA). These membranes were then blocked with 5% milk or 5% BSA and probed with antibodies specific to RBM3 (Abcam Cambridge, United Kingdom), VEGFA (Santacruz Dallas, TX), ZEB-1 (Cell Signaling Technology Denver, MA), SNAI2 (Cell Signaling Technology Denver, MA), TWIST (Cell Signaling Technology Denver, MA) and E Cadherin (Cell signaling Denver, MA). The specific proteins were detected by the enhanced chemiluminescence system (GE Health Care, Piscataway, NJ). Protein expression was captured by Bio-Rad ChemiDoc-XRS+ instrument and densitometric analysis was carried out with Image lab software.

### Immunohistochemistry

Colon cancer tissue microarray, including primary, metastasis and normal tissue, with associated TNM and clinical stage, and pathology grade was purchased from Biomax (Cat. No. CO702b, 69 cases/69 cores). Paraffin-embedded tumor xenografts were subjected to immunohistochemical analysis. Paraffin-embedded tissues were cut to 4μm sections and deparaffinized followed by antigen retrieval. The tissue sections were blocked with the UltraVision Hydrogen Peroxide block for 10 mins (Thermo Scientific). The slides were incubated with primary antibodies for overnight at 4**°**C. were RBM3 (Abcam Abcam Cambridge, United Kingdom), VEGFA (Santacruz), MMP2 (Santacruz Dallas, TX) and E Cadherin (Cell Signaling Technology, Denver, MA). The next day, the primary antibody was washed, and tissues were incubated with HRP Polymer Quanto for 10 mins then developed with a DAB Quanto Chromogen-Substrate mixture. Finally, the slides were counterstained with hematoxylin and eosin. The slides were examined in the Nikon Eclipse Ti microscope under a 20X objective. The TMA slides were scored by pathologist and composite score calculated as (intensity X percent positive cells) along with statistical analysis.

### RNA-sequencing

RNA was extracted from the HCT116 and DLD1 empty vector and RBM3 overexpressing cells. For RNA-immunoprecipitation, colon cancer cells grown to ∼90% confluency in 150-mm dishes were first washed two times with ice-cold phosphate-buffered saline, following which lysates were prepared a polysomal lysis buffer consisting of 100 mmol/L KCl, 25 mmol/L EDTA, 5 mmol/L MgCl2, 10 mmol/L HEPES, pH 7.0, 0.5% Nonidet P-40, 10% glycerol, 2 mmol/L dithiothreitol, 0.4 mmol/L vanadyl ribonucleoside complex, a complete protease inhibitor (Roche Applied Sciences Penzberg, Germany), and RNaseOUT (Invitrogen Carlsbad, CA).

Subsequently, Protein A/G Plus Agarose beads (Thermo Fisher Waltham, MA) were incubated at 4**°**C for at least 18 h with either anti-RBM3 monoclonal antibody (Abcam, Cambridge, United Kingdom) or anti-rabbit-IgG-antibody (Cell Signaling, Danvers, MA). The beads coated with antibody were resuspended in NT2 buffer supplemented with RNaseOUT, 0.2% vanadyl ribonucleoside complex, 2 mmol/L dithiothreitol, and 25 mmol/L EDTA. Supernatant from this lysed material was incubated with the antibody coated beads at 4**°**C with constant mixing for four hr. After incubation, the beads were spun down and washed three times with ice-cold NT2 buffer. Subsequently, the material was digested with proteinase K for 2 hr and RNA extracted with TRIzol reagent.

The RNA quality was analyzed using an Agilent 2100 Bioanalyzer (Agilent Technologies, Santa Clara, CA) according to the manufacturer protocol. RNA sequencing was done by Quick Biology Inc, Pasadena, CA. The samples that passed QC were further used for library preparation. RNA sequencing **l**ibraries were prepared with KAPA Stranded RNA-Seq Kit. The workflow consists of mRNA enrichment, cDNA generation, and end repair to generate blunt ends, A-tailing, adaptor ligation, and PCR amplification. Different adaptors were used for multiplexing samples in one lane. Sequencing was performed on Illumina Hiseq3000 for a pair-end 150 run. Data quality check was done on Illumina SAV. Demultiplexing was performed with Illumina Bcl2fastq 2 v 2.17 program.

The reads were first mapped to the latest UCSC transcript set using Bowtie2 version 2.1.0 and the gene expression level was estimated using RSEM v1.2.15 ^65^. TMM (trimmed mean of M-values) was used to normalize the gene expression. Differentially expressed genes were identified using the edgeR program. Genes showing altered expression with p < 0.05 and more than two-fold changes were considered differentially expressed. The pathway and network analysis were performed using Ingenuity (IPA). IPA computes a score for each network according to the fit of the set of supplied focus genes. These scores indicate the likelihood of focus genes to belong to a network versus those obtained by chance. A score > two indicates a ≤ 99% confidence that a focus gene network was not generated by chance alone. The canonical pathways generated by IPA are the most significant for the uploaded data set. Fischer’s exact test with the FDR option was used to calculate the significance of the canonical pathway.

The lncRNA transcript sets were downloaded from Lncpedia^66^ (https://www.lncipedia.org/). The reads were mapped to the Lncpedia transcript set using Bowtie2 version 2.1.0 ^67^ and the gene expression level was estimated using RSEM v1.2.15. Trimmed mean of M-values (TMM) were used to normalize the gene expression. Differentially expressed genes were identified using the edgeR program ^68^. Genes showing altered expression with p < 0.05 with a more than two-fold change, were considered differentially expressed.

### GO analysis

The Gene Ontology (GO) terms enriched in lncRNA-seq for HCT116 RBM3 overexpressing cells, compared to control having the lowest over-represented *p* values were analyzed by REVIGO (http://revigo.irb.hr) ^30^. The GO terms that are more closely related circles are in closer proximity. The size of the circle indicates the number of mutated genes. The statistical significance of the enriched GO terms is represented by the color of the circle which is based on the over-represented *p*-value. The graph from the revigo was modified for illustration. These GO enrichment terms were also plotted as bar graphs against the p-value.

### Lnc-RNA structure and lncRNA-mRNA interaction prediction

The secondary structure of lncRNA lnc-HOTAIR, lnc-TUG1, lnc-Flii-1, lnc-LSAMP-3 along with the secondary structure of the two IRES in VEGFA mRNA was predicted using RNAfold ^31^ (The ViennaRNA Web Services, http://rna.tbi.univie.ac.at/) and illustrated using VARNA GUI. Along with the prediction of the most stable structure that would be in a state of equilibrium, the software is able to generate minimum free energy information for each sequence. Based on previous published methods, the structure of the lncRNA is colored by the base-pairing probabilities with the unpaired regions colored by the probability of being paired ^31^. Putative interactions between lnc-Flii-1 or lnc-LSAMP-3 with the 5’ and 3’UTRs of VEGFA, ZEB1, SNAI2, and TWIST mRNA were determined using IntaRNA ^32^ (http://rna.informatik.uni-freiburg.de/IntaRNA). Hybridization scores for lncRNA-mRNA interaction were calculated using the IntaRNA tool. The mRNA-lncRNA interactions and RNA secondary structures <u>were modified for illustration.</u>

### Scratch Plate assay

To determine colon cancer cell migration, scratch plate assay was performed in 6-well dishes (Becton Dickinson, Franklin Lakes, NJ). Briefly, 5×10^4^ cancer cells were grown to in 10% serum supplemented DMEM medium. When the culture is nearly confluent, they were incubated overnight in serum-free medium. Scratches were then made in the monolayer and subsequently incubates in 10% serum supplemented DMEM. The plates were photographed at 4X or 10X magnification images every three hr for up to 12 hr in the Cytation3 imaging reader (BioTek, Winooski, VT). The area of the scratch at zero (hr) and 12 (hr) was calculated using ImageJ and percent migration calculated and statistically analyzed.

### Trans-well migration and invasion assay

For studying cancer cell chemotaxis, we used a Boyden chamber which consists of cell culture insert containing 8-μm pore polyethylene terephthalate membrane (EMD Millipore, Burlington, MA). The inserts were seated in 24-well companion plates. For invasion assay, the insert contained Matrigel as a thin gel layer. We seeded 5 x 10^4^ HCT116, 1 x 10^5^ DLD1 or 2 x 10^5^ RKO cells were seeded in the cell culutre insert in serum-free media and placed it in a 24-well plate containing 10% serum containing media. Plates were incubated at 37**°**C with 5% CO2. At the end of the experiment, cells that remained in the upper chamber, which represents nonmigratory cells were removed with a cotton swab. Cells that have moved through the pores and to the bottom of the membrane were fixed with formalin. These were then stained with Hoechst stain and the quantified. For quantification, we chose three fields randomly and photographed them. The experiment was repeated three times.

### Tubule formation assay

Briefly, 15000 HUVEC cells were seeded per well of a 96 well plate on a layer of matrigel (BD Biosciences, Bedford, MA). Cells were treated with condition media from HCT116 or DLD1, empty vector or RBM3 overexpressing cells. After 6 (hr), images of each well were taken under an inverted light microscope. HUVEC cells form a mesh like closed networks of vessel-like tubules. The tube length was calculated using angiogenesis counter in ImageJ software. The experiment was repeated three times and measurements were statistically analyzed and plotted.

### Spheroid assay

Cells, seeded at low density (HCT116, 50 cell/well or DLD1, 100 cells/well) in 96 well low adhesion plates were cultured in DMEM that is supplemented with 20 ng/ml bFGF, 1 mL per 50 mL of 50X B27 supplement and EGF 20 ng/ml (all from Life Technologies, Carlsbad, CA). After 5-7 days, the number of spheroids was determined. To assess the propensity of cells to migrate from spheroids, spheroids were transferred to 96 well attachment plate having serum containing DMEM media. Images of each well were taken on Cytation 3 imaging reader (Biotek, Winooski, VT) from zero to 24 hr at 4 hr intervals. The area of spheroid was calculated at zero hr and 24 hr using ImageJ software. The difference in the area between the 24 and zero hr time points were calculated and the percent migration relative to empty vector cells was determined. The percent migration of RBM3 OE cells relative to the empty vector was calculated and plotted. The experiment was repeated three times and cumulative data was plotted on a bar graph with ± SEM. Make sure to include somewhere how many replicate wells were used per experiment.

### siRNA and LNA gapmer transfection

We designed siRNA for lnc-FLii-1 and lnc-LSAMP-3 using the siRNA Design tool (Dharmacon, Lafayette, CO). These siRNA was purchased along with Lincode non-targeting siRNA#1 control from Dharmacon. The LNA gapmers for lnc-FLii-1 and lnc-LSAMP-3 were designed and purchased along with negative control LNA from Qiagen. For transfection, Lipofectamine-2000 (Invitrogen, Lafayette, CO) and OptiMEM were used according to manufacturer recommendations (Life Technologies). 2 X10^5^ cells were plated in each well of 6 well dish and transfected with a combination of scrambled siRNA and LNA or LSAMP-3 or Flii-1 specific siRNA and LNA. The concentration of siRNA and LNA gapmer used was 200 nM each and mixed with the transfection reagent, Lipofectamine-2000 (Invitrogen, Lafayette, CO). Six hr following transfection, cells were washed with PBS and treated with 10% serum containing DMEM. Twenty-four hr following siRNA transfection, cells were then assayed for migration on the scratch plate assay and RTPCR studies as indicated above. The conditioned medium was collected 24 hr after transfection for tubule formation assay. The experiments were performed at least in triplicates and the experiments repeated three times. All data plotted was statistically analyzed.

### RBM3 and lncRNA truncations

The RBM3 protein was truncated to encompass just the RRM domain and GAS domain. PCR primers were designed encompassing the RRM and GAS domain. The primers included the flag tag and sequence for S6 transcriptase. The lncRNAs were truncated by designing primers encompassing 200-300 bp at the 5’ and 3’ end of each lncRNA. The primer for lncRNA contained the S6 transcriptase. The cDNA from HCT116 cells was used for PCR using the designed primers. The PCR products were run on agarose gel to validate the correct size. The PCR products of appropriate size were purified and further used for in vitro transcription and translation.

### In vitro transcription and translation

The RBM3 and lncRNA truncation PCR products were further utilized for transcription using the MAX1script SP6 transcription kit (#AM1334; Thermo-Fisher, Waltham, MA) following the manufacturer’s protocol with PCR products containing the SP6 promoter. RNA was extracted by phenol/chloroform/isoamyl alcohol (25:24:1) and precipitated with 3 M sodium acetate (pH, 5.2) and 100% ethanol. To label RNA for downstream RNA pull-down assay, RNAs were 3′ end-labeled with a biotinylation kit (#20160; Thermo-Fisher, Waltham, MA) following the manufacturer’s instructions.

### Immunoprecipitation

To identify the domain in RBM3 that interacts with the lncRNAs, lysates from HCT116 cells were prepared with the stably expressing cells using a polysome lysis buffer [100 mmol/L KCl, 25 mmol/L EDTA, 5 mmol/L MgCl2, 10 mmol/L HEPES, pH 7.0, 0.5% Nonidet P-40, 10% glycerol, 2 mmol/L dithiothreitol, 0.4 mmol/L vanadyl ribonucleoside complex, complete protease inhibitor (Roche Applied Sciences Penzberg, Germany), and RNaseOUT (Invitrogen Carlsbad, CA)]. After removal of particulate material by centrifugation, the supernatant was incubated overnight at 4**°**C with the full length and truncated proteins and anti-flag beads in NT2 buffer supplemented with RNaseOUT, 0.2% vanadyl ribonucleoside complex, 2 mmol/L dithiothreitol, and 25 mmol/L EDTA. Subsequently, the beads were spun down, the material was treated with proteinase K, and RNA extracted with TRIzol reagent. The lncRNA interacting with RBM3 truncated protein were identified by PCR using specific primers.

To identify the region on the lncRNA where RBM3 binds, we performed immunoprecipitation using the in vitro transcribed truncated lncRNAs. The truncated lncRNAs were incubated individually with either full length, RRM or GAS domain of RBM3 and were precipitated using anti-flag beads. The lncRNA region interacting with RBM3 truncation was determined by PCR using specific primers for that region.

### Statistical analyses

All statistics calculations were executed using the GraphPad Prism Software (v. 8.1.2, GraphPad Software, San Diego, CA). To determine significance between the groups, Student *t* tests are used. If not specifically stated, the data reported in the manuscript is mean ± standard error of the mean (SEM). For correlation analysis, we used Pearson correlation. Comparison of survival curves was done by Log-Rank (Mantel-cox) test. To assess whether there is significance in tumor volumes and weights, and for biomarker determinations with immunohistochemistry, we performed non-parametric, Mann-Whitney test. Statistical significance between test groups determined by *p* <0.05. All experiments were validated by two or more biological repeats.

## Acknowledgments

**Funding:** This work was supported by Thomas O’Sullivan Foundation, a pilot grant from the NCI-designated University of Kansas Cancer Center (P30CA168524; SA), K-INBRE (NIH-P20 GM103418, PD), BRTP (KUMC, PD) and Schlumberger Foundation (AAAS). S. Anant is an Eminent Scientist of the Kansas Biosciences Authority.

## Author contributions

S.A. conceptualized the research. A.S. performed experiments related to *in silico* prediction and interaction of lncRNA, mRNA, and RBM3. S.G. performed an analysis of sequencing data. A.S. performed *in vitro* experiments. A.S., D.S., and S.C. performed animal experiments. A.S. and S.P. performed western blot analysis. A.S. and P.D. performed IHC staining. O.T. and M.N. evaluated and scored stained IHC slides and COAC TMA slides. A.D., T.I., S.U and R.A.J. contributed to experimental design and interpretation of data. S.A. and S.M.T. supervised the overall research design and data analyses and interpretation. A.S., S.M.T., and S.A. wrote the manuscript. All authors read the manuscript and approved the study.

## 7. Graphical Abstract

We found that RBM3 interacts with mRNA and lncRNA. We also found, using *in silico* analysis that the lncRNA could interact with the 5’UTR either through IRES or IRES independent manner. The lncRNA could interact 5’UTR along with 3’UTR, forming a bridge to keep the mRNA in a circular structure. The working model for our hypothesis is that RBM3 along with the lncRNA can interact with the mRNA 5’UTR and 3’UTR helping and stabilizing the circular structure which enables facilitate multiple rounds of mRNA translation through polysome formation.

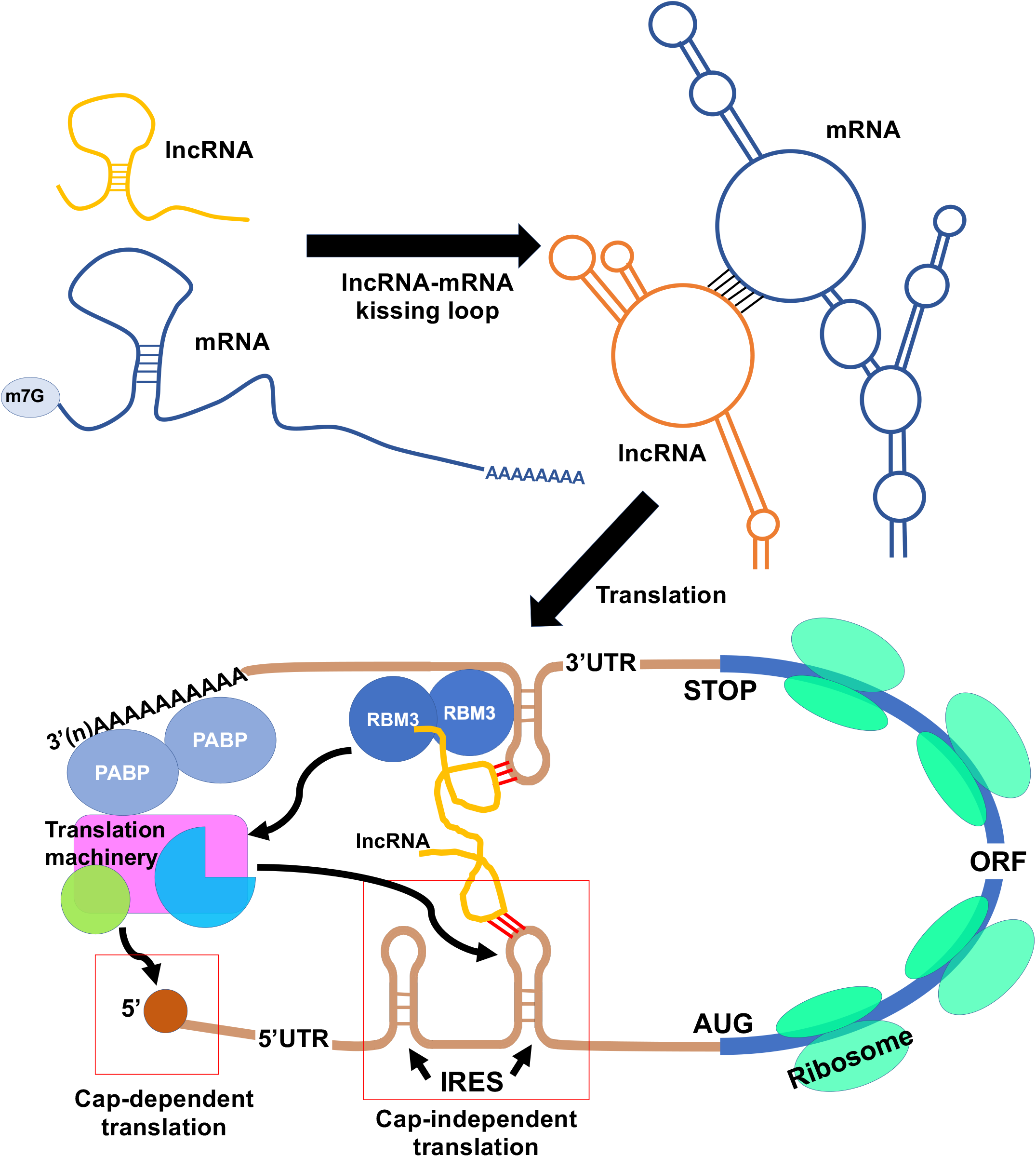

