## Supplementary data for "RNA binding protein RBM3 augments kissing loop formation with lncRNAs to enhance translational control"

**Table 1:** Patient characteristics of cDNA array from Figure.1b showing sex, age, tumor grade, and tumor stage.

**Table 2:** Patient characteristics of tumor microarray [59] from Figure. 1c showing sex, age, tumor grade, tumor stage, and metastasis.

**Table 3A-D:** Table showing predicted the interaction of lnc-LSAMP-3, lnc-Flii-1 lnc-HOTAIR and lnc-TUG1 with the 5'UTR and 3'UTR of VEGFA (A), ZEB1 (B), SNAI2 (C) and TWIST1 (D). The table indicates the position of lncRNA and mRNA interaction along with the minimal energy associated with the interaction.

**Supplementary Figure 1:**

A) RBM3 mRNA levels expressed as transcripts per million in different stages of colon cancer were increased compared to normal ($p<0.001$) as seen in the Cancer Genome Atlas (TCGA) database.
B) Distribution of RBM3 mRNA levels from the TCGA database expressed as transcripts per million in different age groups in colon cancer patients compared to normal.
C) Distribution of RBM3 mRNA levels from the TCGA database expressed as transcripts per million in males and females were increased compared to normal ($p<0.001$) cancer compared to normal.
D) Distribution of RBM3 mRNA levels from the TCGA database expressed as transcripts per million in colon adenocarcinoma and mucinous carcinoma were increased compared to normal ($p<0.001$).

**Supplementary Figure 2:**

A) RBM3 was overexpressed in three colon cancer cell lines (RKO, HCT116, and DLD1) the increased RBM3 expression is demonstrated by western blot analysis.
B) Heat plot depicting the distribution of lncRNA-seq in HCT116 and DLD1 cell lines overexpressing RBM3 compared to empty vector.
C) Predicted Minimum Free Energy (MFE) secondary structure of lncRNA (lnc-HOTAIR, lnc-TUG1, lnc-Flii-1, lnc-LSAMP-3) using RNAfold web server.
D) Quantitative Polymerase Chain Reaction (Q-PCR) show increased levels of lncRNA TUG1 in HCT116 ($p=0.026$) and DLD1 ($p=0.034$) cell lines overexpressing RBM3

compared to empty vector.

E) Correlation analysis between the mRNA expression of RBM3 and lnc-HOTAIR in colon adenocarcinoma obtained using the GEPIA browser. The Pearson Correlation Coefficient R-value is 0.29 and the p-value is 0.56.

F) Correlation analysis between the mRNA expression of RBM3 and lnc-TUG1 in colon adenocarcinoma obtained using the GEPIA browser. The Pearson Correlation Coefficient R-value is 0.12 and p-value is 0.03.

G) The construct used for generating RBM3 overexpressing transgenic mice. The construct was inserted in ROSA26 locus of mice.

**Supplementary Figure 3:**

A) Representative images of the tube formation assay. Increased tube formation is seen in representative images of HUVEC cells treated with condition media from HCT116 and DLD1 RBM3 overexpressing cells, compared to empty vector cells.

B) Increase in the number of spheroids in RBM3 overexpressed HCT116 and DLD1 cells as seen in representative images.

C) Plot for number of spheroids shows significant increase in the number of spheroids in RBM3 overexpressed HCT116 (p=0.047) and DLD1 (p<0.01) cells compared to the empty vector cells.

D) Representative images of the spread of migratory cells from the spheroid from Fig. 4c at time zero (hr) and 24 (hr). Images from HCT116 and DLD1 empty vector and RBM3 overexpressing cells are shown.

E) Plot for percentage spread of migratory cells from the spheroid shows increased spread of HCT116 (p=0.012) and DLD1 (p=0.02) RBM3 overexpressing cells compared to empty vector.

F) Representative images of scratch plate assay of HCT116, DLD1 or RKO empty vector and RBM3 overexpressing cells from Figure. 4d. Images at time 0hr and 12 (hr) are shown.

G) Representative images of transwell migration assay of HCT116, DLD1 or RKO empty vector and RBM3 overexpressing cells from Figure. 4e. Images at time zero (hr) and 12 (hr) are shown.

H) Representative images of Transwell invasion assay of HCT116, DLD1 or RKO empty vector and RBM3 overexpressing cells from Figure. 4d. Images at time zero (hr) and 12 (hr) are shown.

**Supplementary Figure 4:**

A) Visualization of mRNA -lncRNA interaction between VEGFA 5'UTR and 3'UTR and lnc-LSAMP-3 and lnc-Flii-1as predicted using IntaRNA.
B) Visualization of mRNA -lncRNA interaction between ZEB1 5'UTR and 3'UTR and lnc-LSAMP-3 and lnc-Flii-1 as predicted using IntaRNA.
C) Visualization of mRNA -lncRNA interaction between SNAil2 5'UTR and 3'UTR and lnc-LSAMP-3 and lnc-Flii-1 as predicted using IntaRNA.
D) Visualization of mRNA -lncRNA interaction between TWIST1 5'UTR and 3'UTR and lnc-LSAMP-3 and lnc-Flii-1 as predicted using IntaRNA.
E) Visualization of mRNA -lncRNA interaction between VEGFA 5'UTR and 3'UTR and lnc-HOTAIR and lnc-TUG1 as predicted using IntaRNA.
F) Visualization of mRNA -lncRNA interaction between ZEB1 5'UTR and 3'UTR and lnc-HOTAIR and lnc-TUG1 as predicted using IntaRNA.
G) Visualization of mRNA -lncRNA interaction between SNAil2 5'UTR and 3'UTR and lnc-HOTAIR and lnc-TUG1 as predicted using IntaRNA.
H) Visualization of mRNA -lncRNA interaction between TWIST1 5'UTR and 3'UTR and lnc-HOTAIR and lnc-TUG1 as predicted using IntaRNA.
I) Predicted Minimum Free Energy (MFE) secondary structure of VEGFA IRES-A in 5'UTR the interactions between VEGFA IRES-A and lnc-HOTAIR, lnc-TUG1, and RBM3 are highlighted.
J) Correlation of the genes and lncRNA with RBM3 in colorectal cancer from TCGA database.

**Supplementary Figure 5:**

A) Immunohistochemistry analysis shows increased composite score for RBM3 levels in HCT116 ($p=0.001$) and DLD1 ($p<0.001$) RBM3 overexpressing xenograft as compared to empty vector.
B) Immunohistochemistry analysis shows increased expression of CD31 levels in HCT116

and DLD1 RBM3 overexpressing xenograft as compared to empty vector.

**Supplementary Figure 6:**

A) Representative images for colony formation of scramble and shRNA knocked down clones of RBM3.

B) The no of colonies formed by the RBM3 shRNA knockdown clones over scramble for (p=0.009), sh3 (p=0.006), sh5 (p=0.0004).

C) The size of colonies formed by the RBM3 shRNA knockdown clones over scramble for (p=0.0007), sh3 (p=0.023), sh5 (p=0.004).

D) Representative images of Transwell migration and invasion assay of HCT116 scramble and shRNA RBM3 clones Figure. 6D. Images at time zero (hr) and 12 (hr) are shown.

E) RT-PCR validation of lncRNA lnc-LSAMP-3 and lnc-Flii-1 in the tumor xenograft tissues from HCT116 scramble and shRNA RBM3 knockdown clone.

F) Schematic for the RBM3 knockout mice generation. The exons 2-6 were chosen for knockout and were floxed. The floxed RBM3 mice, these mice crossed with mice carrying a colon-specific promoter driven Cre (CDX2-driven-Cre/ERT2) construct from Jackson Labs for colon specific knockout. The conditional knockout for RBM3 in the colon was done by tamoxifen treatment to mice containing the RBM3 floxed gene and the CDX2-Cre.

G) Representative gel image of genotyping RBM3 floxed mice. Wt band 410bp and floxed band 523bp

H) Representative gel image of PCR for RBM3 after RBM3 kockout.

I) Schematic for the timeline of the AOM/DSS induced carcinogenesis model on the Cre inducible RBM3 knockout mice.

J) Representative images of the distal colon with tumors after the AOM/DSS induced carcinogenesis in RBM3 knockout and wild type mice.

K) Representative images of H and E staining the distal colon with tumors after the AOM/DSS induced carcinogenesis in RBM3 knockout and wild type mice. Marked the area of adenocarcinoma.

L) Plot for change in the tumor size after the AOM/DSS induced carcinogenesis in RBM3 knockout and wild type mice.

**Supplementary Figure 7:**

(A) Representative images of the endothelial tubular network of HUVEC cell treated with condition media from lncRNA knockdown in HCT116 empty vector and RBM3 overexpressing cell. The images of three wells were taken and the experiment repeated three times.

(B) Schematic of xenograft model. Colon cancer cell lines overexpressing RBM3 and empty vector ($1 \times 10^6$) cells were inoculated subcutaneously into the right flank of athymic Foxn1nu mice. The lncRNA knockdown was performed by intratumoral injection of si+LNA gapmer (10uM each time) for each lncRNA.

Supplementary Figure 1

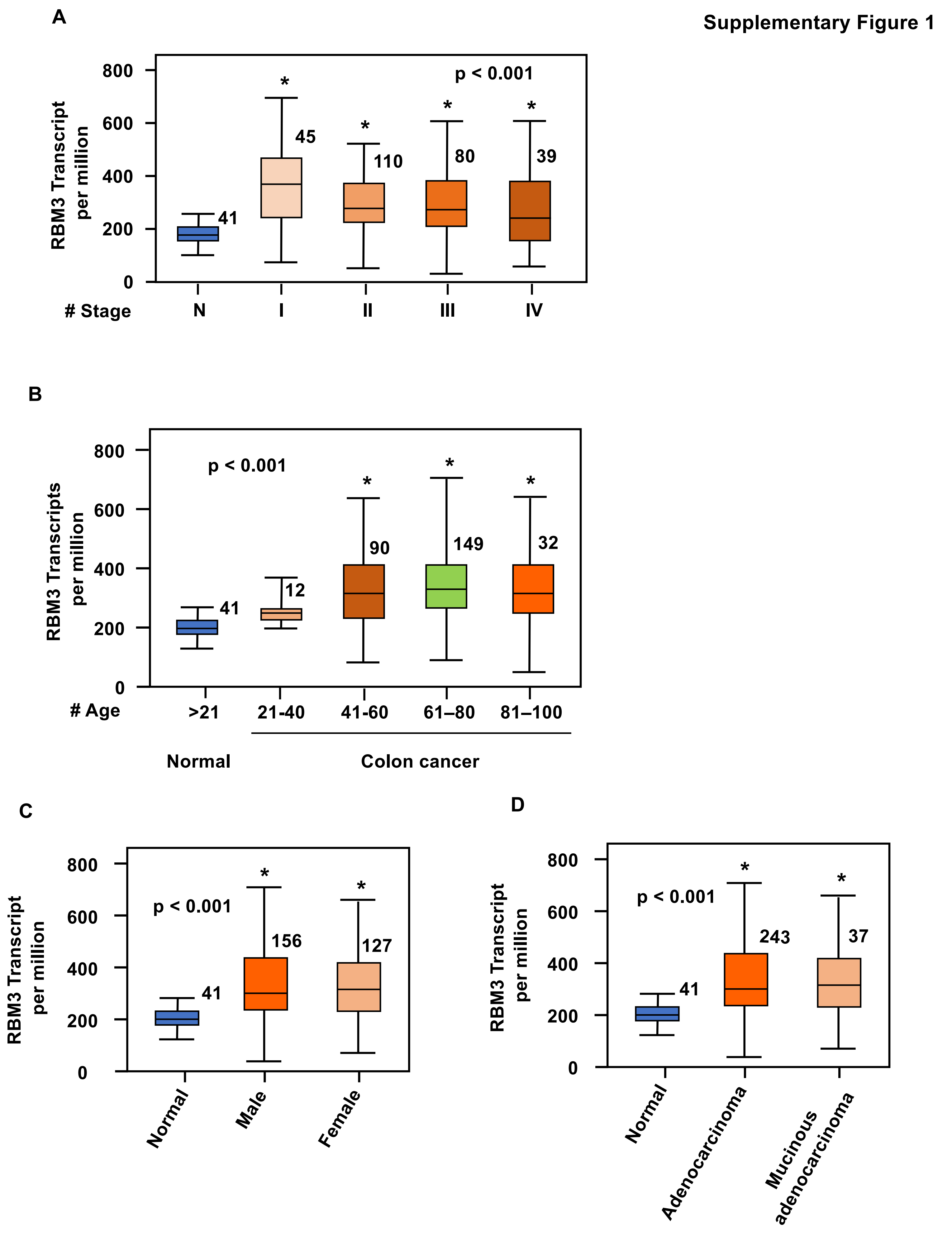

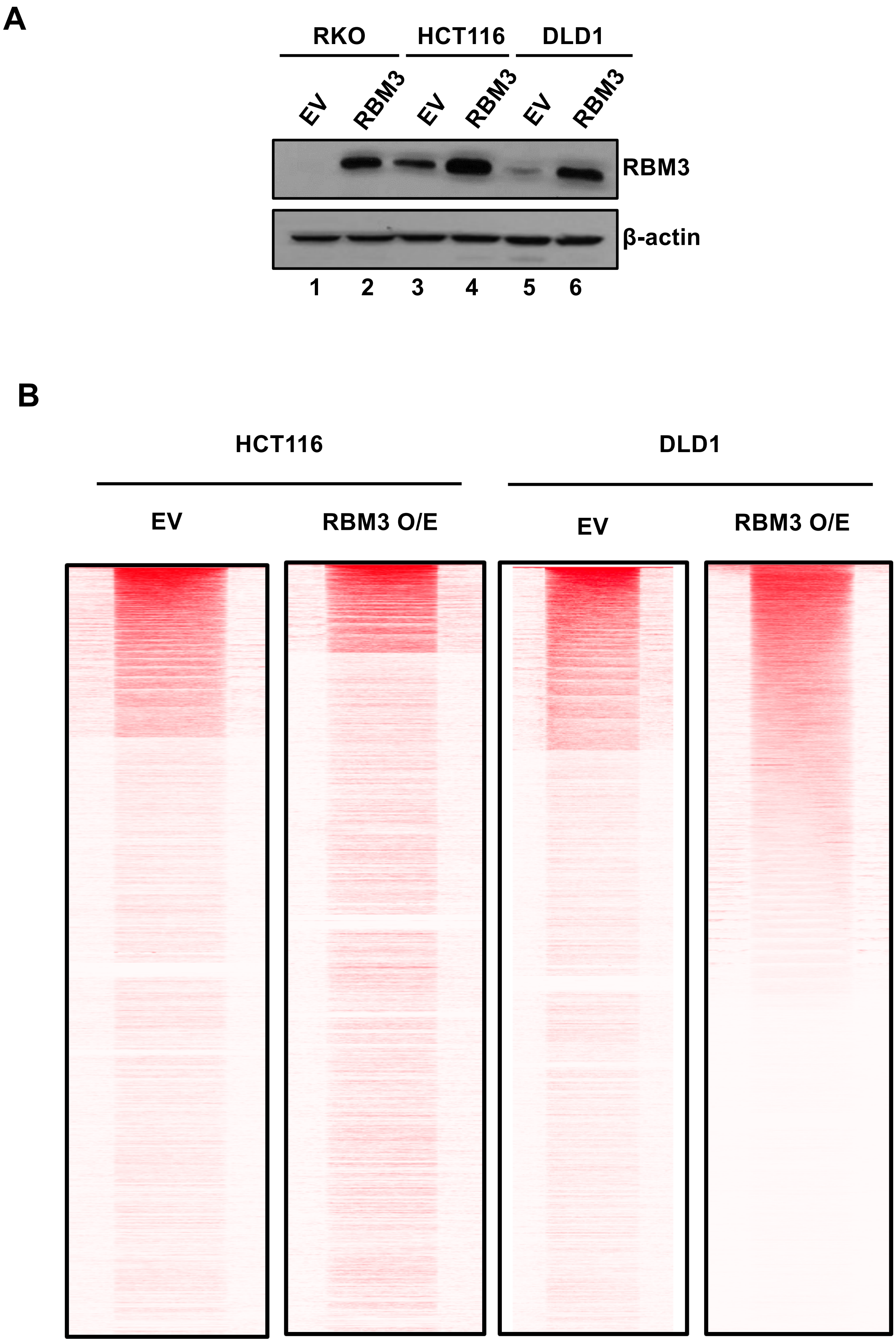
A
RKO
HCT116
DLD1
EV
RBM3
EV
RBM3
EV
RBM3
RBM3
β-actin
1
2
3
4
5
6
B
HCT116
DLD1
EV
RBM3 O/E
EV
RBM3 O/E

**Supplementary Figure 2**

**C**

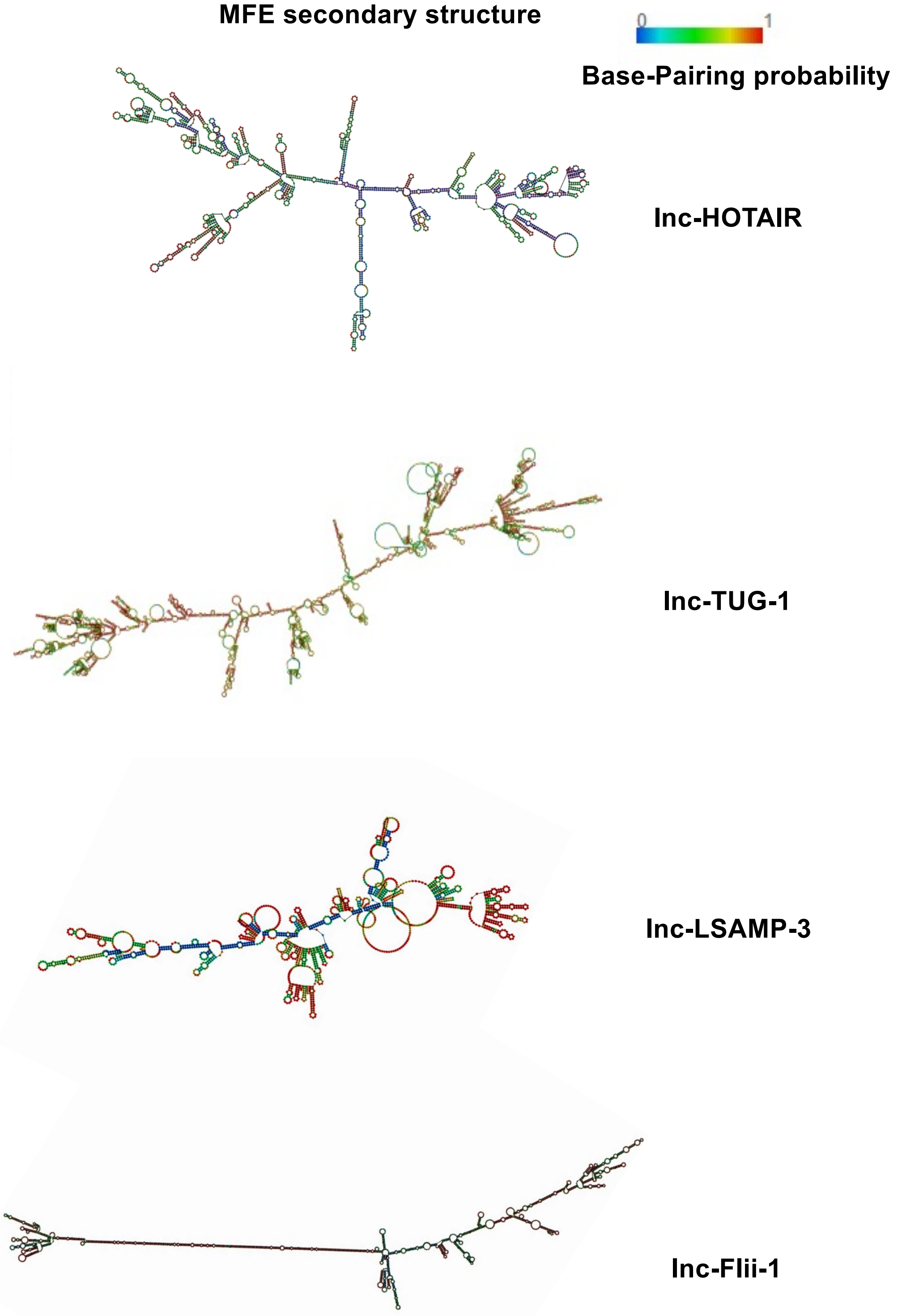

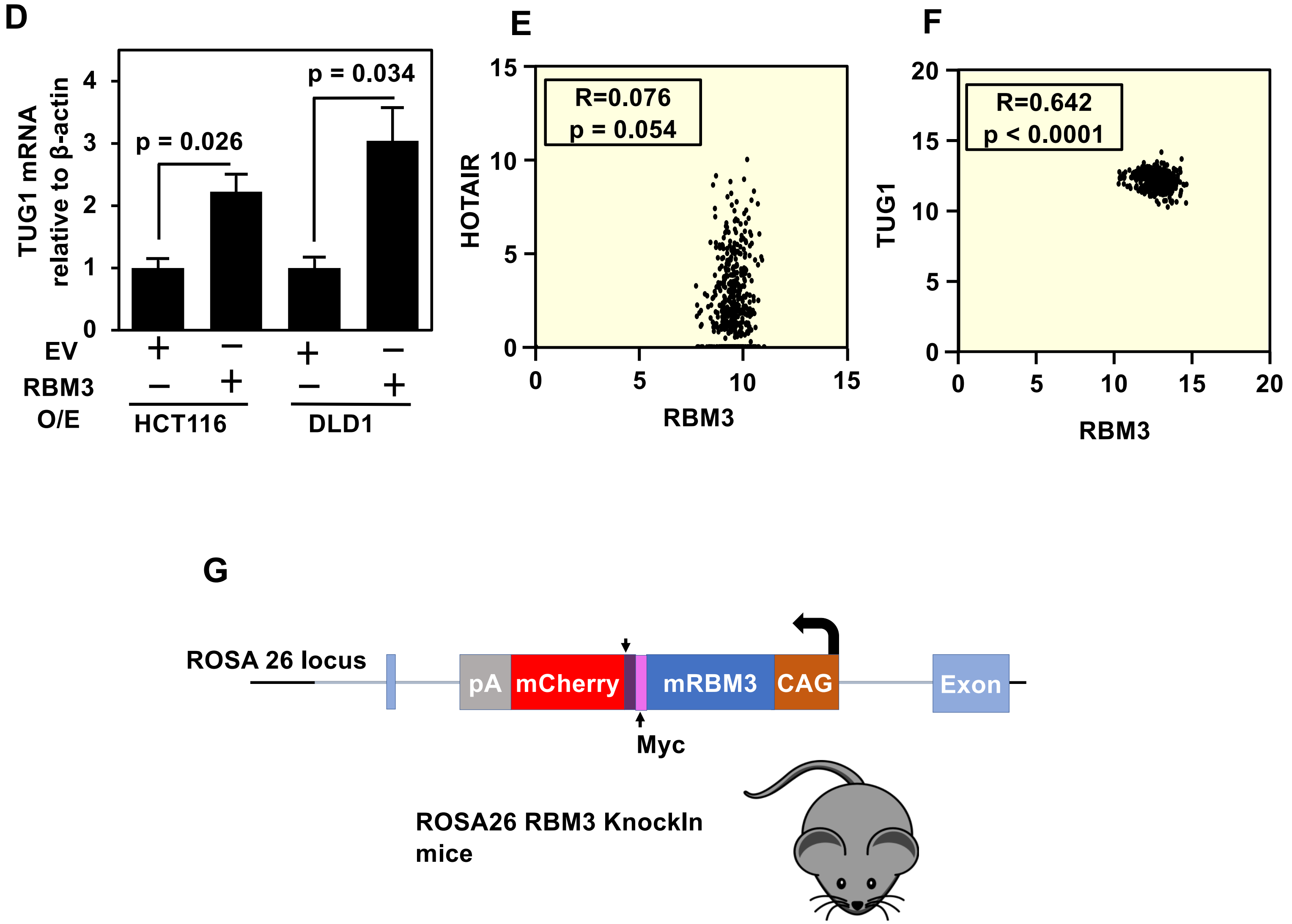
D
TUG1 mRNA
relative to β-actin
p = 0.026
p = 0.034
4
3
2
1
0
EV
RBM3
O/E
HCT116
DLD1
E
R=0.076
p = 0.054
HOTAIR
RBM3
15
10
5
0
F
R=0.642
p < 0.0001
TUG1
RBM3
20
15
10
5
0
G
ROSA 26 locus
pA
mCherry
mRBM3
CAG
Exon
Myc
ROSA26 RBM3 KnockIn
mice

**Supplementary Figure 3**

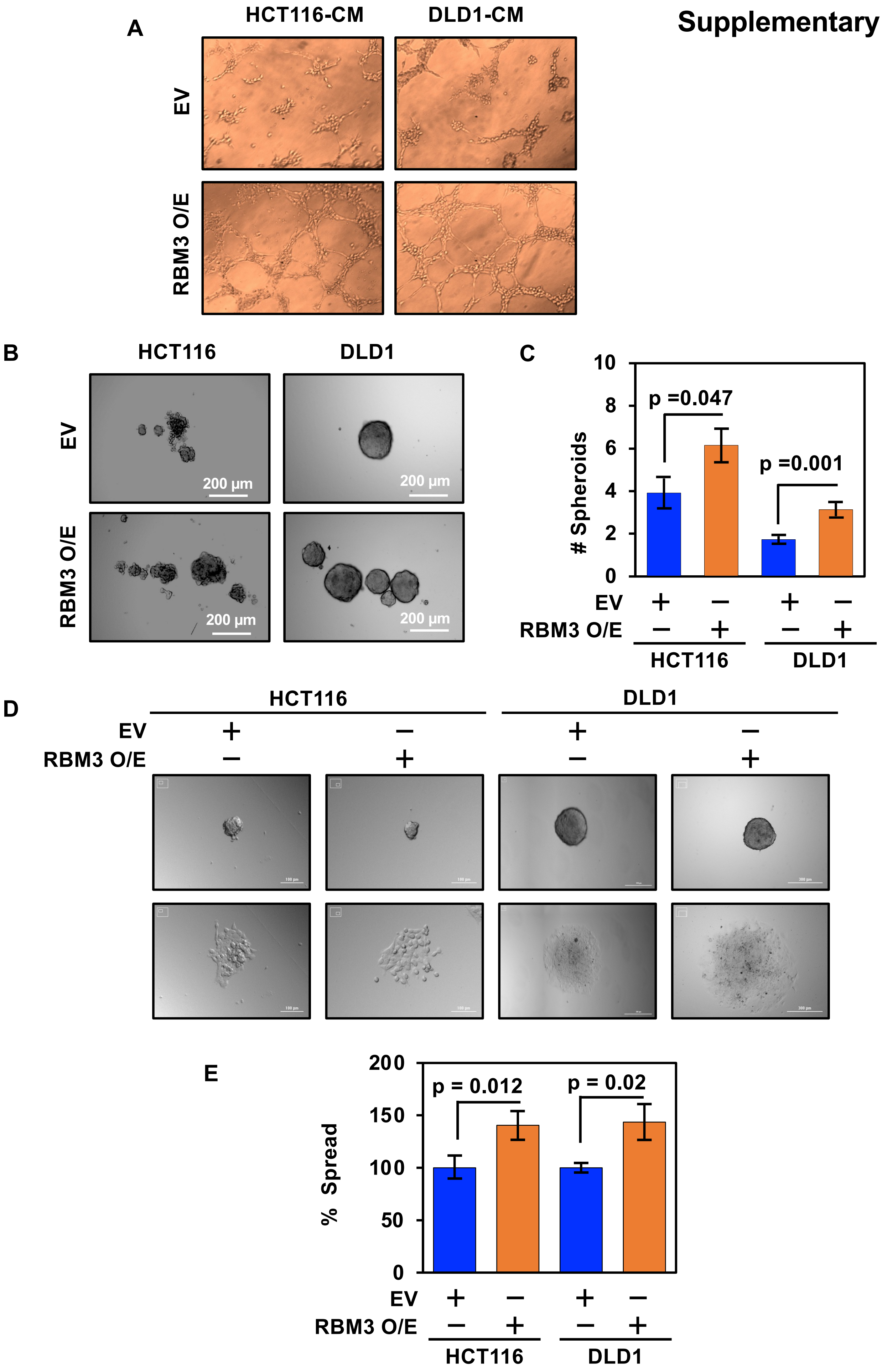

### Supplementary Figure 3

F

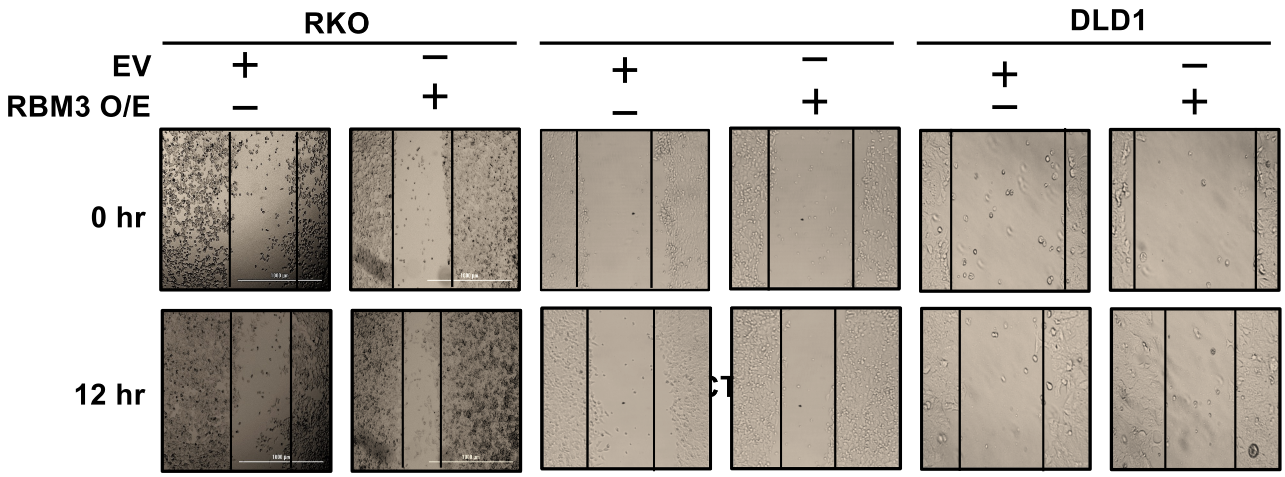


G

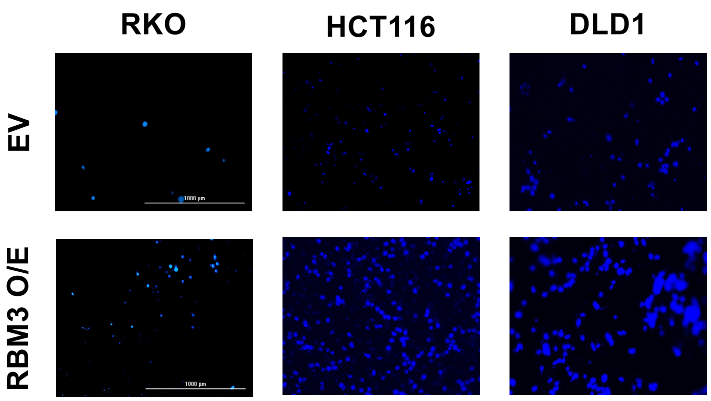


H

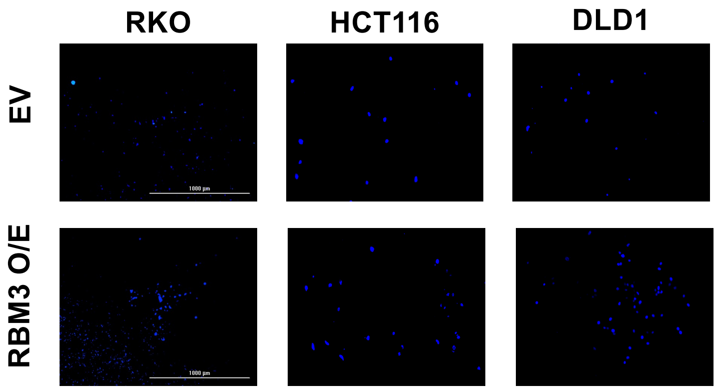

**Supplementary Figure 4**

**A**

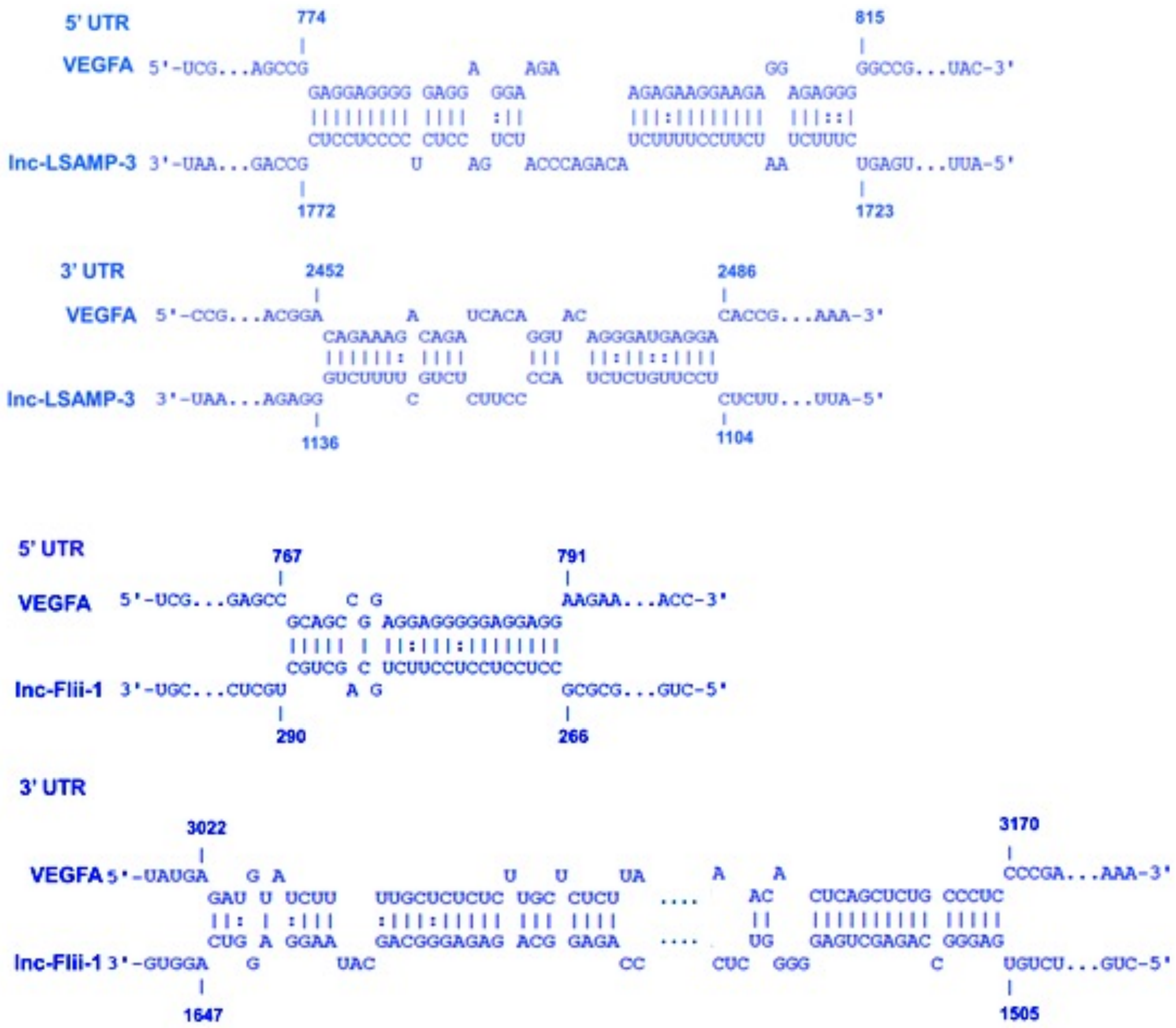

**Supplementary Figure 4**

**B**

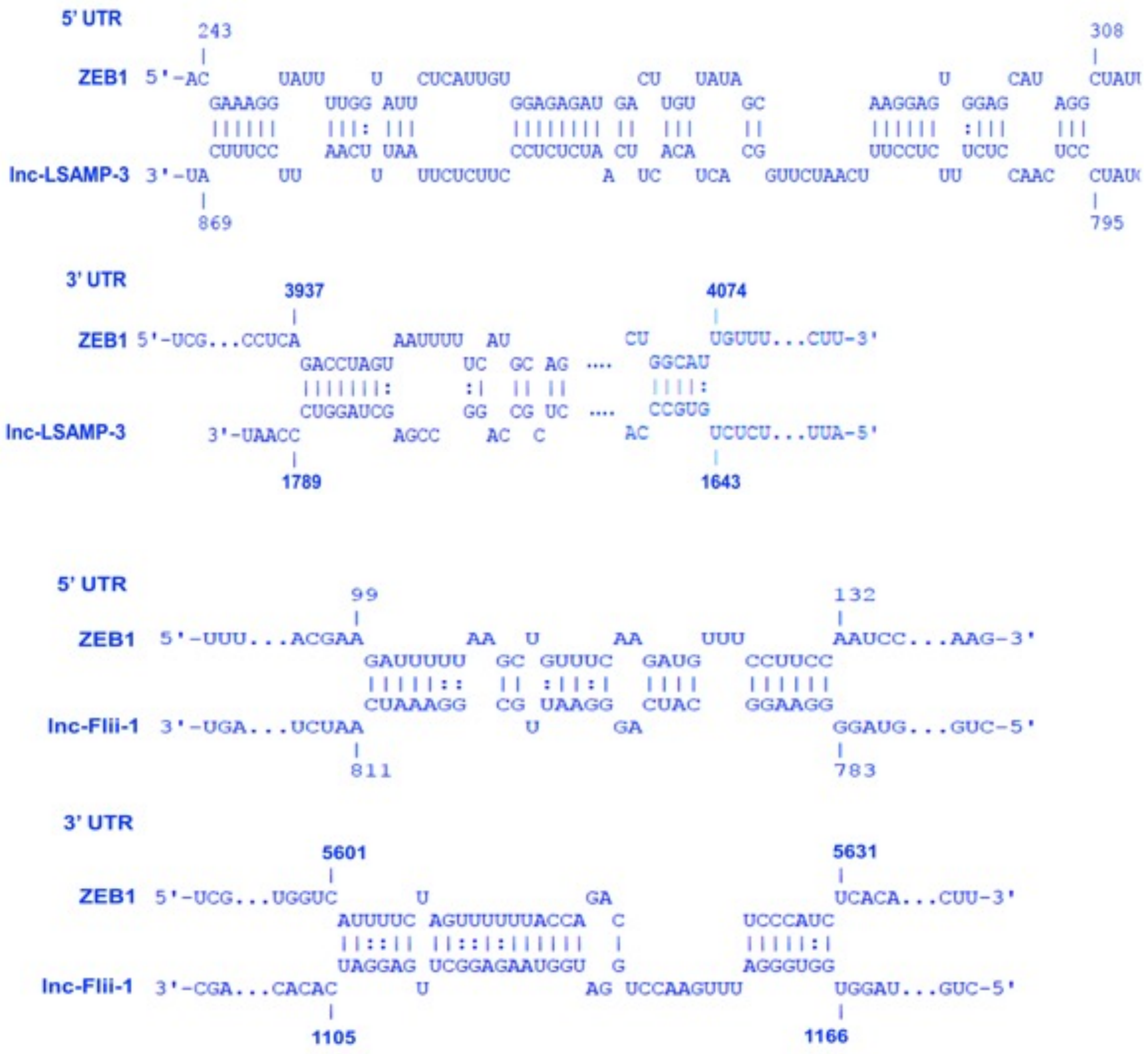

**Supplementary Figure 4**

**C**

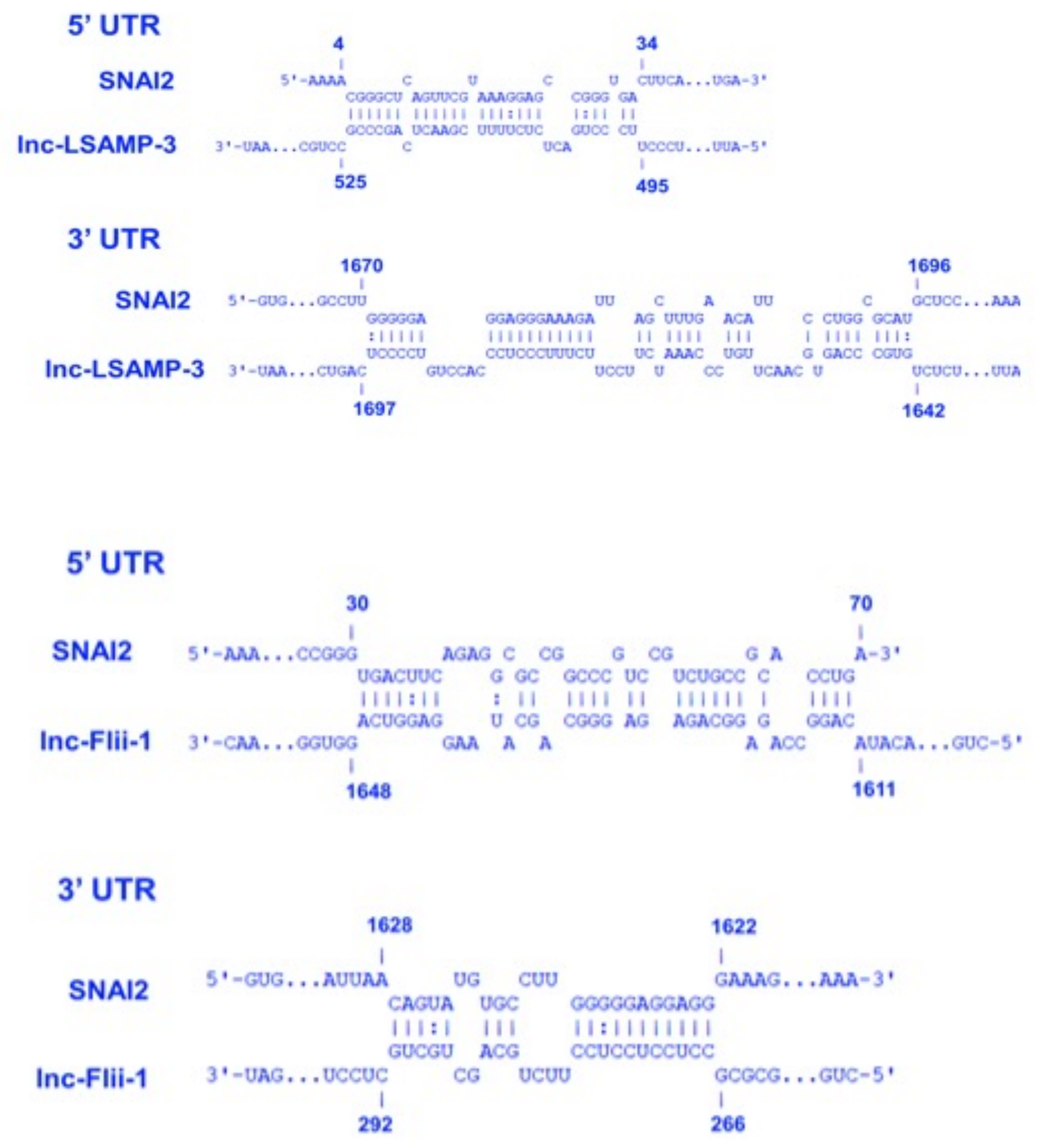

Supplementary Figure 4

D

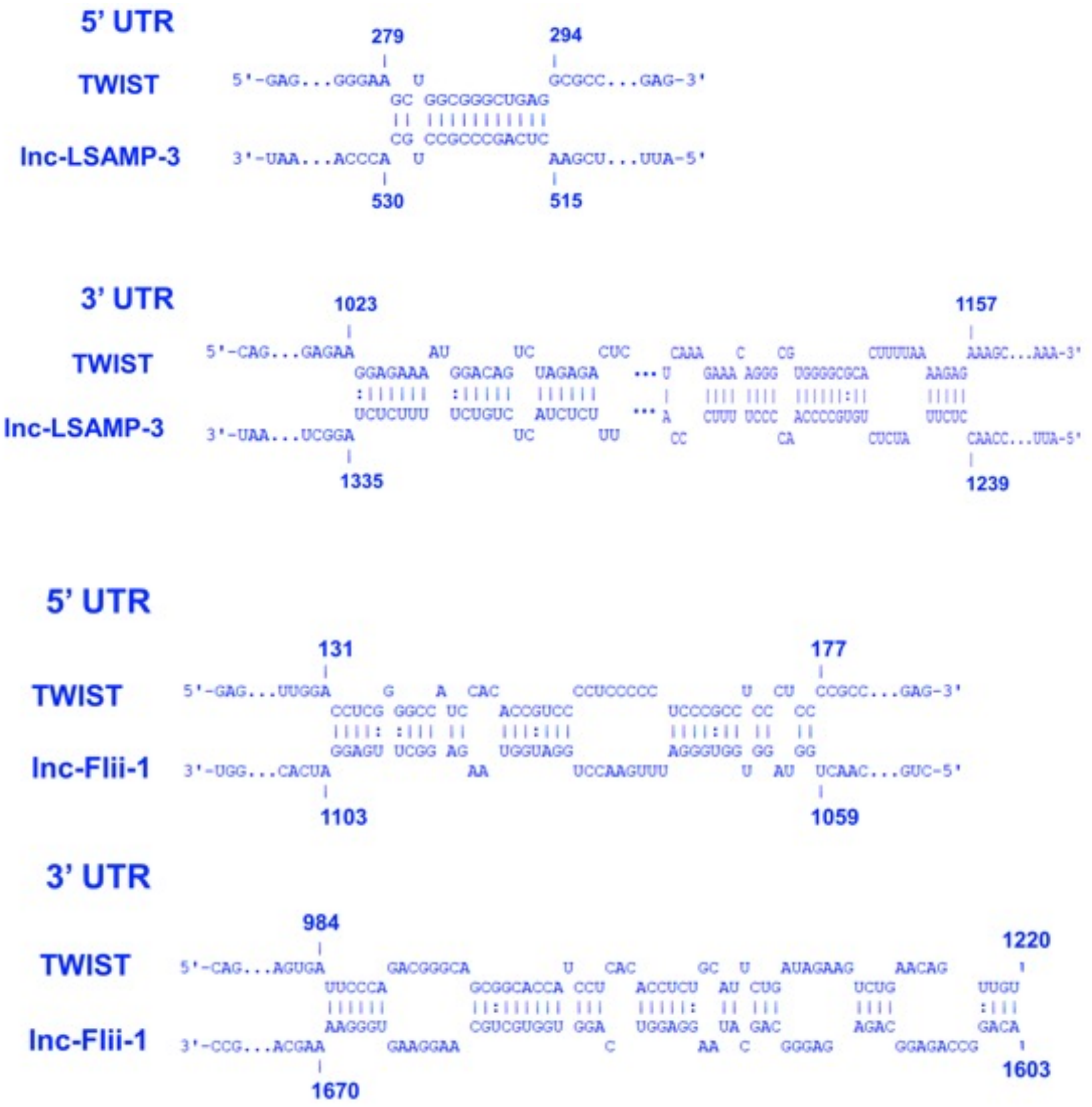

**E**

**Supplementary Figure 4**

**5'UTR**

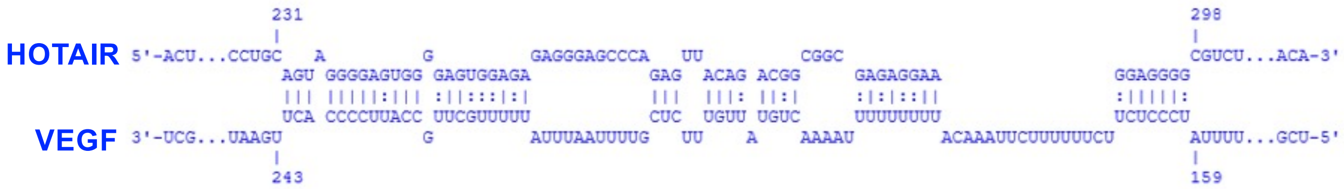


**3'UTR**

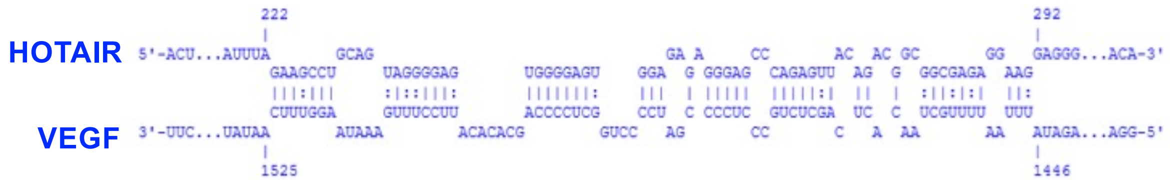


**5'UTR**

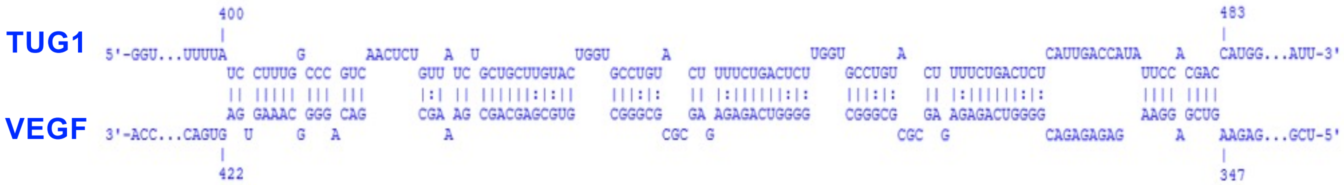


**3'UTR**

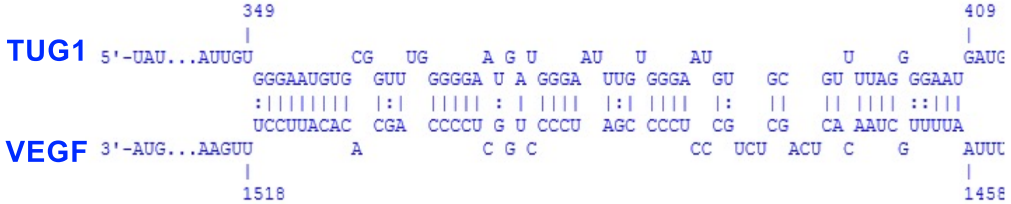

**F** **Supplementary Figure 4**

**5’UTR**

```
               298            310
                 |              |
ZEB1    5'-GUG..UUGAG  G        UGCUG..UUU-3'
                 CU UUGCCGCUGU
                 || ||||||||||
                 GA AGCGGCGGCA
HOTAIR  3'-AGA..GGAAG  G        GACAU..ACC-5'
                 |              |
               427            415
```

**3’UTR**

```
             1717                              1748
               |                                 |
ZEB1    5'-AAU..AAUCA    AACC          AU     A    GCCAA..CAU-3'
               GCCU    AUACAACUC  UC  UUCCUU GUAA
               ||||    |||||||||  ||  |||||| ||||
               CGGA    UGUGUUGGG  AG  AAGGAA CAUU
HOTAIR  3'-CAA..AAGGC              GA  CG          AAGAA..ACC-5'
               |                                 |
             1443                              1415
```

**5’UTR**

```
               295                        316
                 |                          |
ZEB1    5'-GUG..AUAUU    U                 UGUGGCUUU-3'
                 GAGC GUUGCC GCUGUUGCUGA
                 |||| |||||| |||||||||||
                 CUCG CGACGG CGGCGGCGGCU
TUG1    3'-CAA..CCCGC    U      G          GGUGG..GGC-5'
                 |                          |
               460                        438
```

**3’UTR**

```
               654            666
                 |              |
ZEB1    5'-AAU..UUAAG           GAAGA..UGG-3'
                 AAAGCAGCUGAUA
                 |||||||||||||
                 UUUCGUCGACUGU
TUG1    3'-ACG..ACUAU           UGAUU..GGC-5'
                 |              |
              1872            1860
```

**G** **Supplementary Figure 4**

**5'UTR**

**SNA2** / **HOTAIR**

```
           37                    53
             |                    |
5'-AGU..CGCCG           G         ACCUG..CCG-3'
             GCCCGU    CC UCUGCCGC
             ||||||    || ||||||||
             CGGGCA    GG AGACGGUG
3'-AGA..ACUCA      GAAC G         CAAAC..ACC-5'
             |                    |
          728                     708
```

**3'UTR**

**HOTAIR** / **SNAI2**

```
          817                                      849
             |                                     |
5'-UCA..AAUAA    AU        CAAGUG                  UAUAG..UUA-3'
             UACA  ACCCCUUC      CCU      UUUUAAAUUG
             ||||  ||||||||      |||      ||||||||||
             AUGU  UGGGGAAG      GGA      AGAAUUUAAC
3'-CAA..GCCGG    GU        CGAA     ACAUUA         CCGAC..ACC-5'
             |                                     |
         1440                                      1404
```

**5'UTR**

**TUG1** / **SNAI2**

```
          479                                                                                 545
             |                                                                                |
5'-CGG..GGGAG    C        CA   G           UU   AG    C                       CG              AGGCC..CUA-3'
             GCGG    GGCGG  GCG CCG CG GGAU   GG  CGGC GG GGA            GUGGC  GGGCC GGCUUGG
             ||||    |||||  ||| ||| || ||||   ||  |||| || |||            |||||  ||||| |||||||
             CGCC    CCGCC  CGC GGC GC CCUA   CC  GCCG CC CCU            CGCCG  CCCGG CCGAGCC
3'-GCC..ACGGU    CAGA          G   C  G    UUG  AG    C  U   GGAUGUCGUAG        U         CGUCC..UGA-5'
             |                                                                                |
          157                                                                                 73
```

**3'UTR**

**SNAI2** / **TUG1**

```
          636                                              681
             |                                            |
5'-UCA..UGCCU   G    G    AA  AUU   UUU   CAUUCCU          CCAUU..UUA-3'
             UGG GGAG AGGG  AG   AGC   GAA        GGCGCAUGCU
             ||| |||| ||||  ||   |||   |||        ||||||||||
             ACC CCUC UCCC  UC   UCG   CUU        CCGCGUGCGA
3'-AGU..AAUUU   G         AG  CUU   UC    CUCU             CAGUG..GGC-5'
             |                                            |
           79                                              39
```

**5’UTR**

```
                  91                                                                                                     160
                     |                                                                                                   |
TWIST1  5'-AAC..UCUUU         C       AUCCACAC        U                                           C      CUC      GCG    CCCCG..GAU-3'
                     UUGGAC CU  GGGGCC               CG CCCCUCC  CC CUC CCGC                CUCCCUC   CCGC   CCCCGC   CCCU
                     |||||| ||  ||||||               || |||||||  || ||| ||||                |||||||   ||||   ||||||   ||||
                     AGUCUG GA  CCCCGG               GC GGGGAGG  GG GAG GGCG                GAGGGAG   GGUG   GGGGUG   GGGA
HOTAIR  3'-CAA..UUUGU      A  AA      AAUUUUUUUAUUUCU          AA  A   C    GCAGACAUUGAGACCC       AGA    A        AG     UGACG..ACC-5'
                     |                                                                                                   |
                 471                                                                                                     375
```

**3’UTR**

```
                  148                                              198
                     |                                            |
TWIST1  5'-CUA..AUUAG      AAAAUCCAAA       AAAC     G    GGCG           GAGAA..GAA-3'
                     AAGAGC          UUCAAAG    AGGGC UGG     CACUUUUAAAA
                     ||||||          |||||||    ||||| |||     |||||||||||
                     UUCUCG          AGGUUUC    UCCCG ACC     GUGAAAAUUUU
HOTAIR  3'-CAA..GGAAG      A                CGA           AAA            UAAAC..ACC-5'
                     |                                            |
                 1608                                              1570
```

**5’UTR**

```
                  104                                      137
                     |                                     |
TWIST1  5'-AAC..CGGGG   U     C          U         C       UCCCC..GAU-3'
                     CCA   CCA ACCGUCC CC  CCCCCUCC GCCUCCC
                     |||   ||| ||||||| ||  |||||||| |||||||
                     GGU   GGU UGGUAGG GG  GGGGGAGG CGGGGGG
TUG1    3'-CAC..UUUAG   GUU           A  CC        A       GUUGA..GGC-5'
                     |                                     |
                  623                                      587
```

**3’UTR**

```
                   13                      30
                     |                     |
TWIST1  5'-CUA..GCGGA      CA              AGGGC..GAA-3'
                     GCCCCC      CCCCCUCAGC
                     ||||||      ||||||||||
                     CGGGGG      GGGGGGGUUG
TUG1    3'-CAA..GAGGC      AGGAC           ACCCA..GGC-5'
                     |                     |
                  603                      583
```

I

Supplementary Figure 4

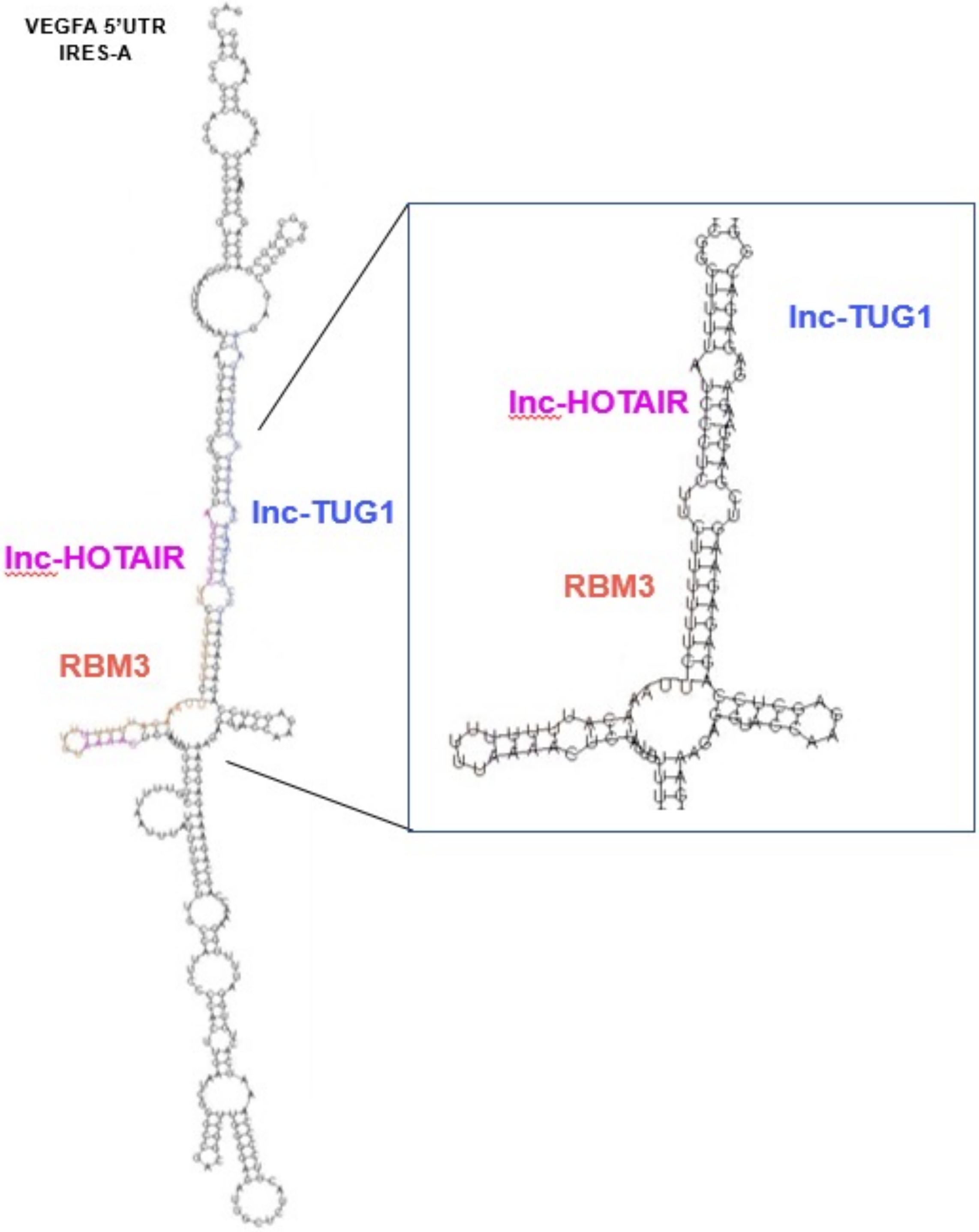

### Supplementary Figure 5

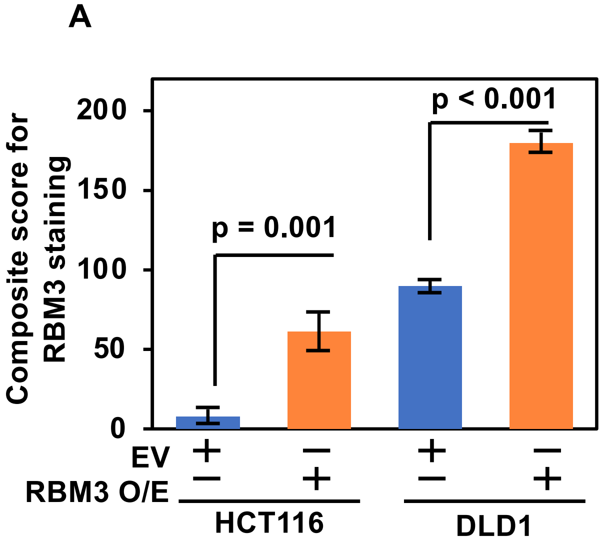


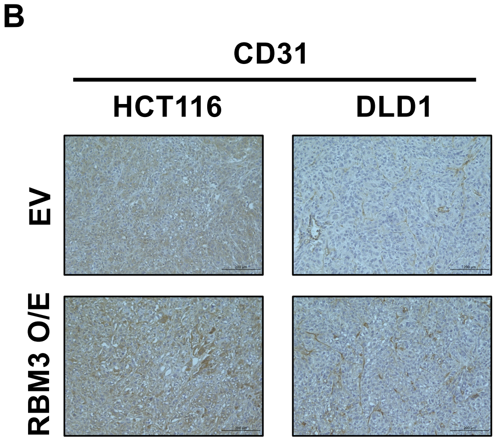

### Supplementary Figure 6

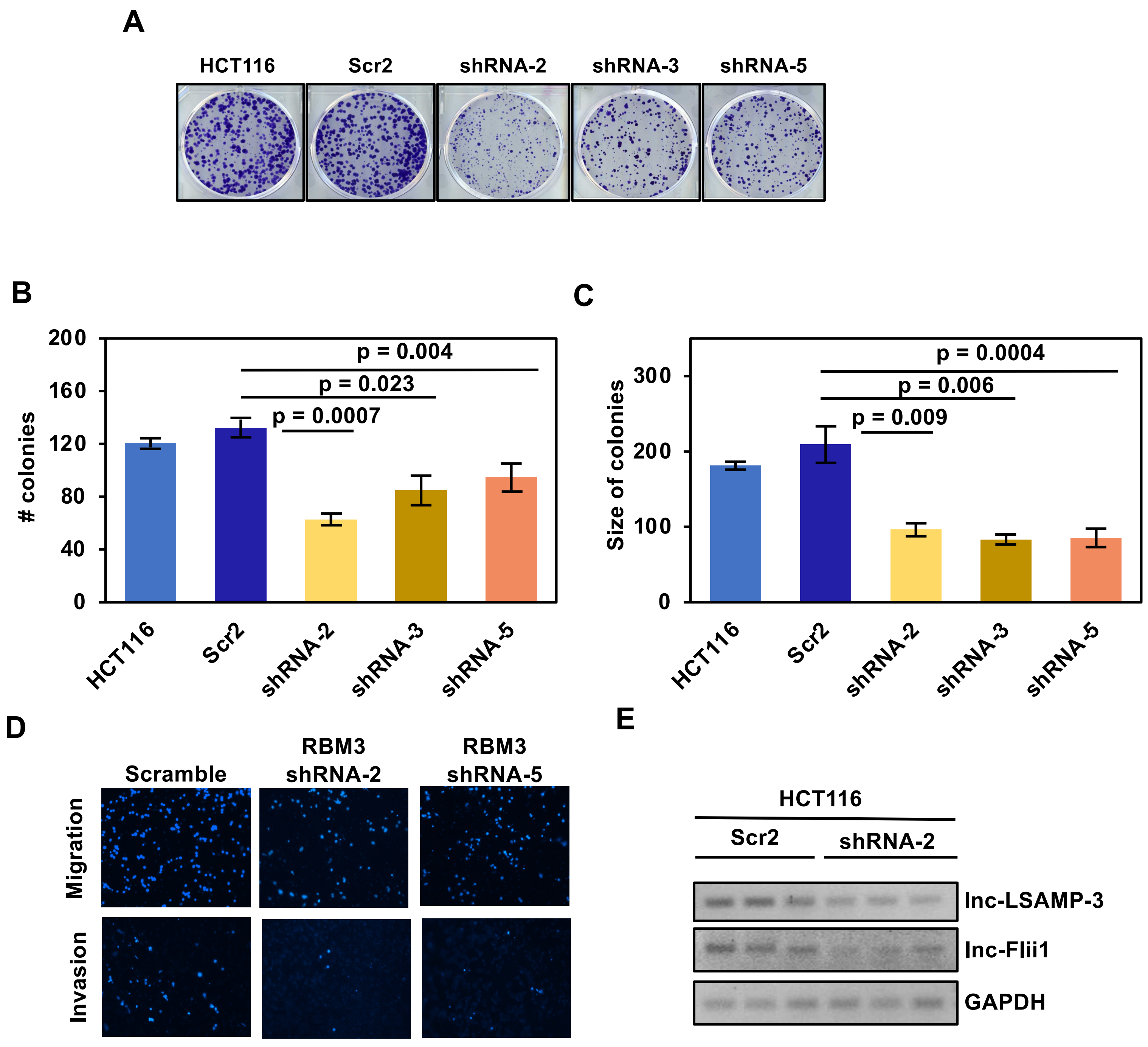

#### Supplementary Figure 6

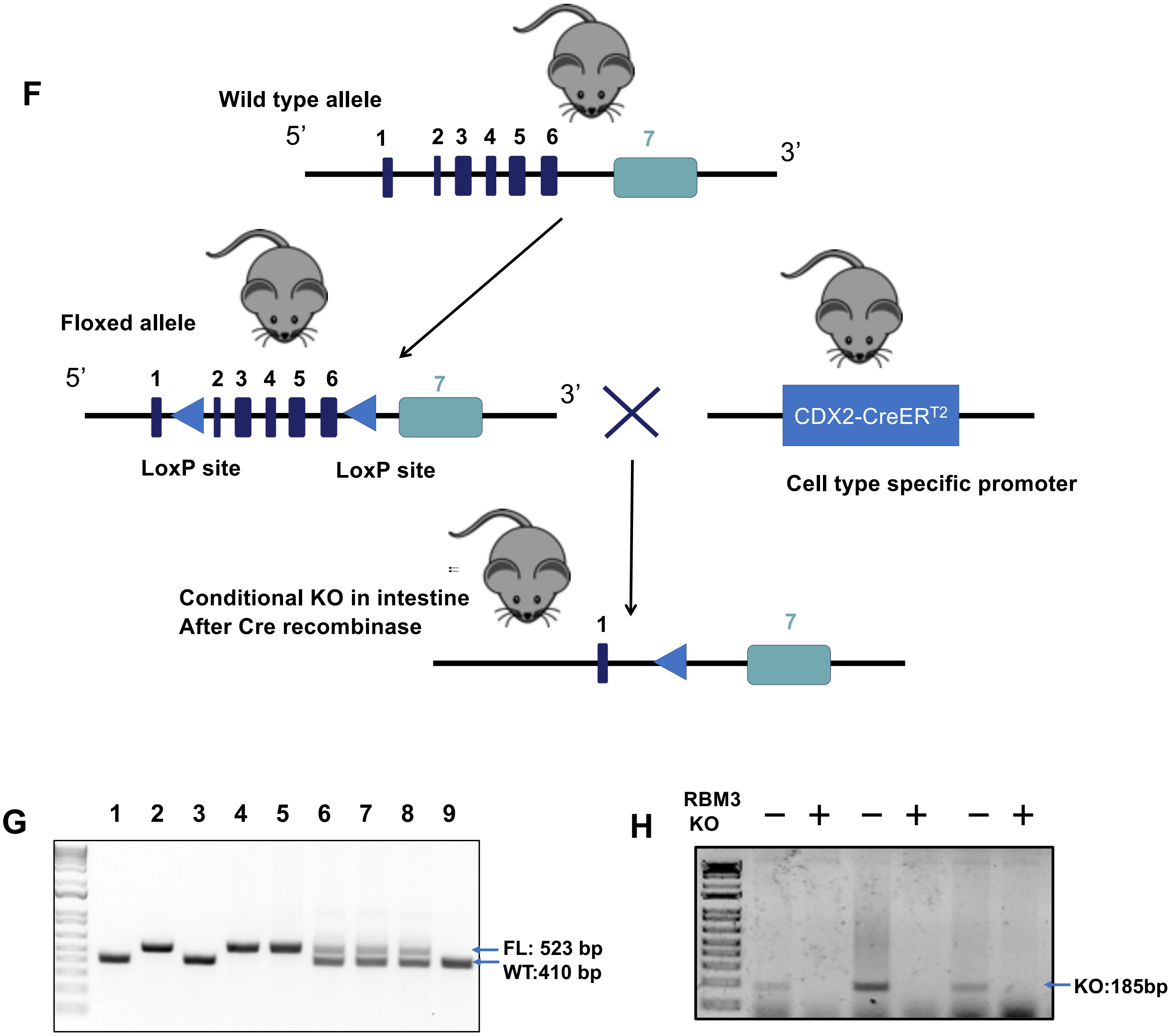

#### Supplementary Figure 6

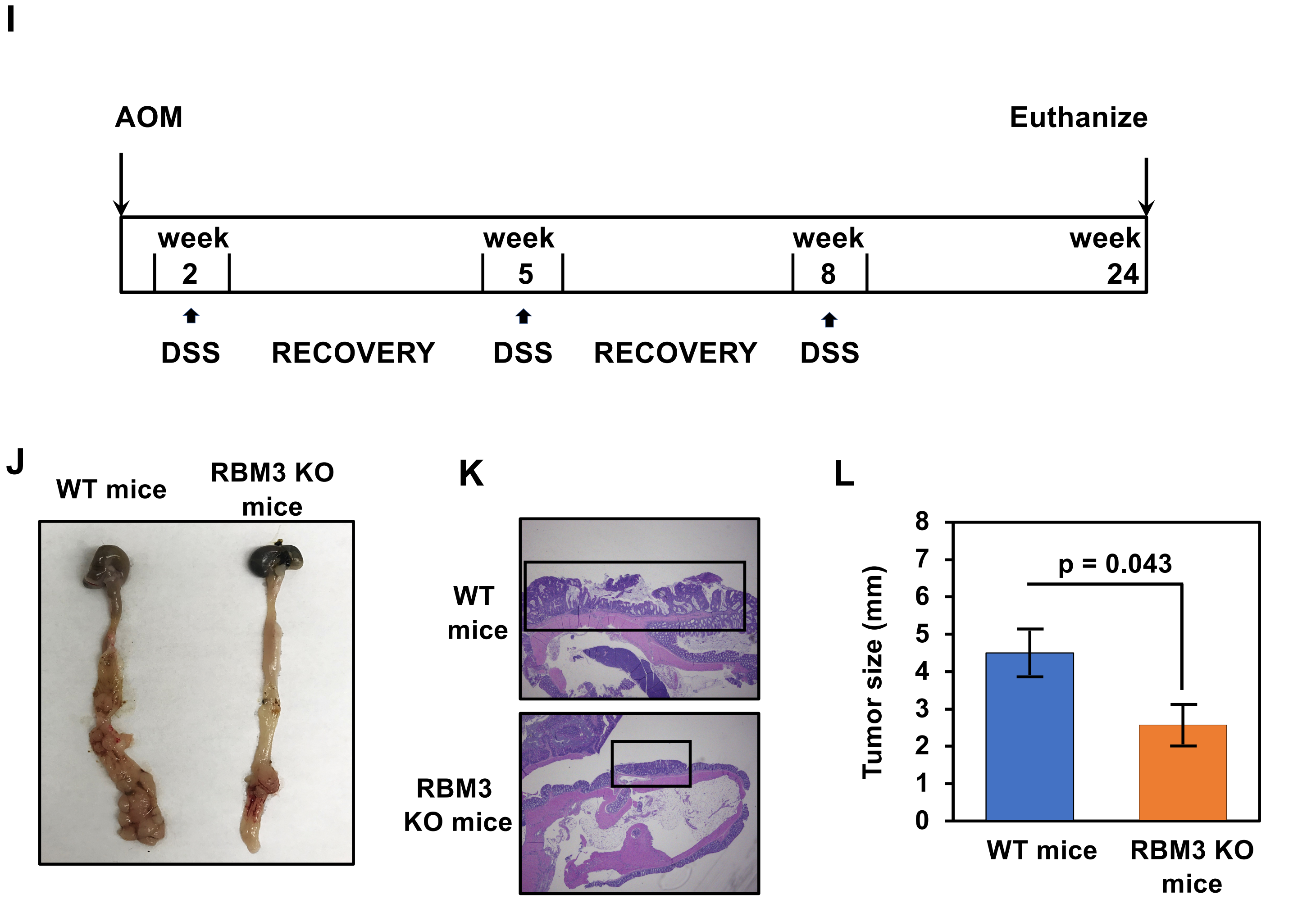

Supplementary Figure 7

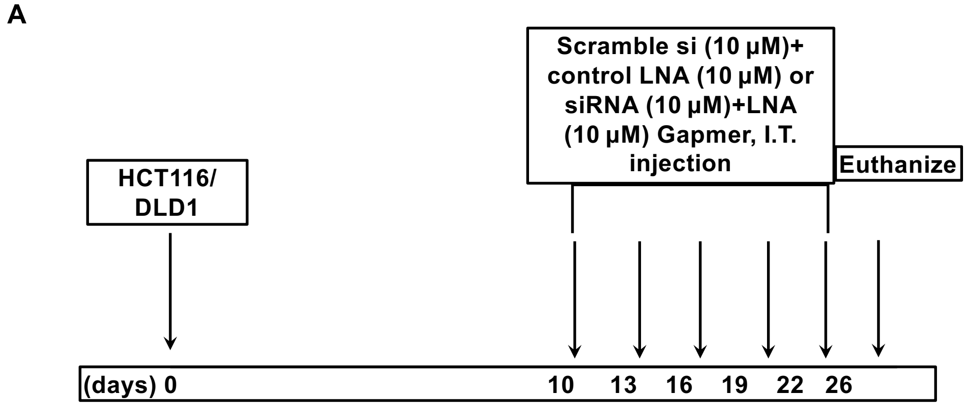


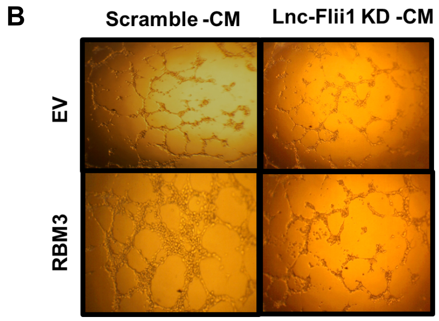

**Table 1**

| Characteristic | | Number (N= 24) |
|---|---|---|
| Sex | Male | 11 |
| | Female | 13 |
| Age | Range | 21-89 |
| | Median | 76 |
| Tumor grade | Moderately differentiated | 15 |
| | Well differentiated | 8 |
| | Poorly differentiated | 1 |
| Tumor stage | I | 5 |
| | II | 7 |
| | III | 8 |
| | IV | 4 |

**Table 2**

| TMA Characteristic | | Number (N=70) | P value |
|---|---|---|---|
| Sex | Male | 51 | 0.730 |
| | Female | 19 | |
| Age | range | Below 52 | 0.693 |
| | median | Above 52 | |
| Tumor grade | Normal | 10 | |
| | Moderately differentiated | 12 | 0.041 |
| | Well differentiated | 22 | 0.051 |
| | Poorly differentiated | 21 | 0.018 |
| Tumor stage | I/II | **15** | 0.017 |
| | III/IV | 12 | 0.041 |
| Metastatic | | 30 | 0.036 |
| | liver | 5 | |
| | lymph node | 25 | |

#### Supplementary Table 3

##### Table A

| | VEGFA | | | | | |
|---|---|---|---|---|---|---|
| | IRES/5'UTR | | | 3'UTR | | |
| lncRNA | mRNA 5'UTR | lncRNA position | DG (Kcal/mol) | mRNA 3'UTR | lncRNA position | DG (Kcal/mol) |
| lnc-LSAMP-3 | 774 – 815 | 1723 – 1772 | -45.2 | 2452 – 2486 | 1104 – 1136 | -20.4 |
| lnc-Flii-1 | 767 – 791 | 290 – 266 | -18.8 | 3022 – 3170 | 1505 – 1647 | -31.5 |
| lnc-HOTAIR | 160 -- 242 | 232 -- 297 | -23.53050 | 1447 -- 1524 | 223 -- 291 | -32.39040 |
| lnc-TUG1 | 401 -- 482 | 348 -- 421 | -34.55190 | 1459 -- 1517 | 350 -- 408 | -23.74930 |

##### Table B

| | ZEB1 | | | | | |
|---|---|---|---|---|---|---|
| | IRES/5'UTR | | | 3'UTR | | |
| lncRNA | mRNA 5'UTR | lncRNA position | DG (Kcal/mol) | mRNA 3'UTR | lncRNA position | DG (Kcal/mol) |
| lnc-LSAMP-3 | 243 – 308 | 795 – 869 | -15.4 | 3937 – 4074 | 1643 – 1789 | -18.7 |
| lnc-Flii-1 | 99 – 132 | 783 – 811 | -12.6 | 5601 – 5631 | 1066 – 1105 | -17.3 |
| lnc-HOTAIR | 298-310 | 415-427 | -13.04 | 1717-1748 | 1415- 1443 | -14.19 |
| lnc-TUG1 | 295-316 | 438-460 | -13.63 | 654-666 | 1860-1872 | -13.08 |

##### Table C

| | SNAI2 | | | | | |
|---|---|---|---|---|---|---|
| | IRES/5'UTR | | | 3'UTR | | |
| lncRNA | mRNA 5'UTR | lncRNA position | DG (Kcal/mol) | mRNA 3'UTR | lncRNA position | DG (Kcal/mol) |
| lnc-LSAMP-3 | 4 – 34 | 495 – 525 | -14.9 | 1670 – 1696 | 1642 – 1697 | -21.8 |
| lnc-Flii-1 | 30 – 70 | 1611 – 1648 | -12.8 | 1628 – 1622 | 266 – 292 | -21.4 |
| lnc-HOTAIR | 37-53 | 708-728 | -18.15 | 817-849 | 1404-1440 | -17.25 |
| lnc-TUG1 | 73-157 | 479-545 | -18.2 | 636-681 | 39-79 | -21.73 |

#### Supplementary Table 3

#### Table D

| | TWIST1 | | | | | |
|---|---|---|---|---|---|---|
| | IRES/5'UTR | | | 3'UTR | | |
| lncRNA | mRNA 5'UTR | lncRNA position | DG (Kcal/mol) | mRNA 3'UTR | lncRNA position | DG (Kcal/mol) |
| lnc-LSAMP-3 | 279 - 294 | 515 – 530 | -18.8 | 1023 – 1157 | 1239 – 1335 | -22.9 |
| lnc-Flii-1 | 132 – 176 | 1060 – 1102 | -21.2 | 984 – 1220 | 1604 – 1671 | -18.9 |
| lnc-HOTAIR | 91-160 | 375-471 | -36.77 | 148-198 | 1570-1608 | -16.91 |
| lnc-TUG1 | 104-137 | 587-623 | -40.68 | 13-30 | 583-603 | -18.37 |
